# An autoinhibitory regulatory switch governs heterotypic phase separation and liquid-to-solid phase transition of TDP-43 and tau into cytotoxic amyloids

**DOI:** 10.64898/2026.05.23.727391

**Authors:** Anuja Walimbe, Snehasis Sarkar, Roopali Khanna, Arpita Singh, Deepti D. Ramteke, Ashish Joshi, Samrat Mukhopadhyay

**Affiliations:** Centre for Protein Science, Design and Engineering, Indian Institute of Science Education and Research (IISER) Mohali, Punjab 140306, India; Department of Biological Sciences, Indian Institute of Science Education and Research (IISER) Mohali, Punjab 140306, India; Department of Chemical Sciences, Indian Institute of Science Education and Research (IISER) Mohali, Punjab 140306, India

## Abstract

Biomolecular condensates formed via macromolecular phase separation of proteins and nucleic acids control a myriad of essential cellular processes, whereas abnormal phase transitions into solid-like aggregates are associated with a range of fatal neurodegenerative diseases. Here, we present a unique case to demonstrate that two neuronal proteins, TDP-43 and tau, undergo heterotypic phase separation via domain-specific interactions regulated by an autoinhibitory conformational switch. Using single-molecule FRET (Förster resonance energy transfer), in combination with multi-color high-resolution imaging, fluorescence recovery after photobleaching, fluorescence correlation spectroscopy, single-droplet fluorescence anisotropy imaging, homoFRET microscopy, fluorescence lifetime-FRET imaging, vibrational Raman spectroscopy, and electron microscopy, we unmask the interplay of molecular drivers and dissect the sequence of events associated with the formation of TDP-43:tau co-condensates that undergo liquid-to-solid phase transitions into cytotoxic amyloids. Our cellular studies show that a disease-associated cytosolic fragment of TDP-43 recruits tau into oxidative stress-induced cytoplasmic granules, eliciting cellular toxicity. Our findings provide mechanistic underpinnings of co-condensation-mediated aberrant phase transitions associated with exacerbated neuropathological outcomes.

## Introduction

A diverse array of cellular functions is orchestrated in a highly spatiotemporally-controlled manner by a wide range of membraneless biomolecular condensates that are formed via macromolecular phase separation of proteins and nucleic acids^1–12^. These biomolecular condensates are highly dynamic, liquid-like, reversible, tunable, non-stoichiometric, and selectively permeable mesoscopic supramolecular assemblies possessing emergent material properties. The formation, maintenance, and disassembly of these intracellular condensates and their physical properties are critically associated with a myriad of fundamental cellular processes, including genome organization, gene regulation, RNA biogenesis, processing, and transport, signal transduction, stress response, immune response, and so forth^4–7, 12–17^. Intrinsically disordered proteins/regions (IDPs/IDRs) possessing low sequence complexity have been recognized as the principal candidates due to their flexibility, structural heterogeneity, and multivalency that together govern the formation of dynamic physical networks via a multitude of transient noncovalent interactions involving electrostatic, hydrophobic, cation-π, and π-π contacts^18–22^. These sequence-encoded interactions can facilitate both homotypic and heterotypic interactions in the complex cellular milieu and drive phase separation of a range of proteins and nucleic acids into multicomponent biomolecular condensates with emergent physical properties^23–26^. Such dynamic liquid-like condensates can undergo irreversible phase transitions into solid-like aggregates that are pathological hallmarks of fatal neurodegenerative diseases, including Alzheimer’s disease (AD), amyotrophic lateral sclerosis (ALS), frontotemporal lobar degeneration (FTLD), and so forth^27–35^.

A central player in the pathology of ALS and FTLD is TDP-43 (TAR DNA-binding protein 43), an RNA-binding protein that plays critical roles in pre-mRNA splicing, mRNA stability, and miRNA biogenesis^31,36,37^. TDP-43 is a 414 amino acid long protein, with a multi-domain architecture comprising a structured N-terminal domain (NTD, residues 1-80) exhibiting a ubiquitin-like fold, two RNA-recognition motifs (RRM1 and RRM2, residues 104-262), followed by a glycine-rich disordered C-terminal domain (CTD, residues 270-414) (Fig. 1a and Supplementary Fig. 1a). Primarily, following its synthesis in the cytoplasm, TDP-43 localizes to the nucleus owing to the bipartite nuclear localization signal (NLS, residues 82-98) responsible for its critical role in RNA metabolism^36,37^. However, cellular conditions such as oxidative, osmotic, or heat stress can lead to the redistribution of TDP-43 from the nucleus to the cytoplasm, and facilitate its recruitment into phase-separated stress granules that sequester translationally stalled mRNAs and other RNA-binding proteins (RBPs)^36–38^. Aberrant phase transitions of such TDP-43-rich granules due to prolonged stress or mislocalization of TDP-43 into the cytoplasm can lead to the formation of irreversible, hyperphosphorylated, solid deposits, associated with ALS and FTLD pathology^39–43^. Such liquid-to-solid phase transitions are further facilitated by abnormal posttranslational modifications and impaired nucleocytoplasmic shuttling due to a point mutation in the NLS or an abnormal cleavage into C-terminal fragments, namely, TDP-35 and TDP-25. An emerging body of evidence has revealed a co-deposition-mediated pathological synergism between TDP-43 and tau, a primarily cytoplasmic IDP that binds microtubules, thereby stabilizing the cellular cytoskeleton. The full-length tau protein (2N4R) is intrinsically disordered with distinct domains, comprising two N-terminal acidic regions (N1 and N2; residues 1-151), a central proline-rich domain (residues 153-244), a repeat region consisting of 5 repeats (R1-R4 and R’; residues 244-390) containing a microtubule-binding region (MTBR; residues 244-372), and a C-terminal domain (Fig. 1b and Supplementary Fig. 1b)^44^.

**Figure 1:**
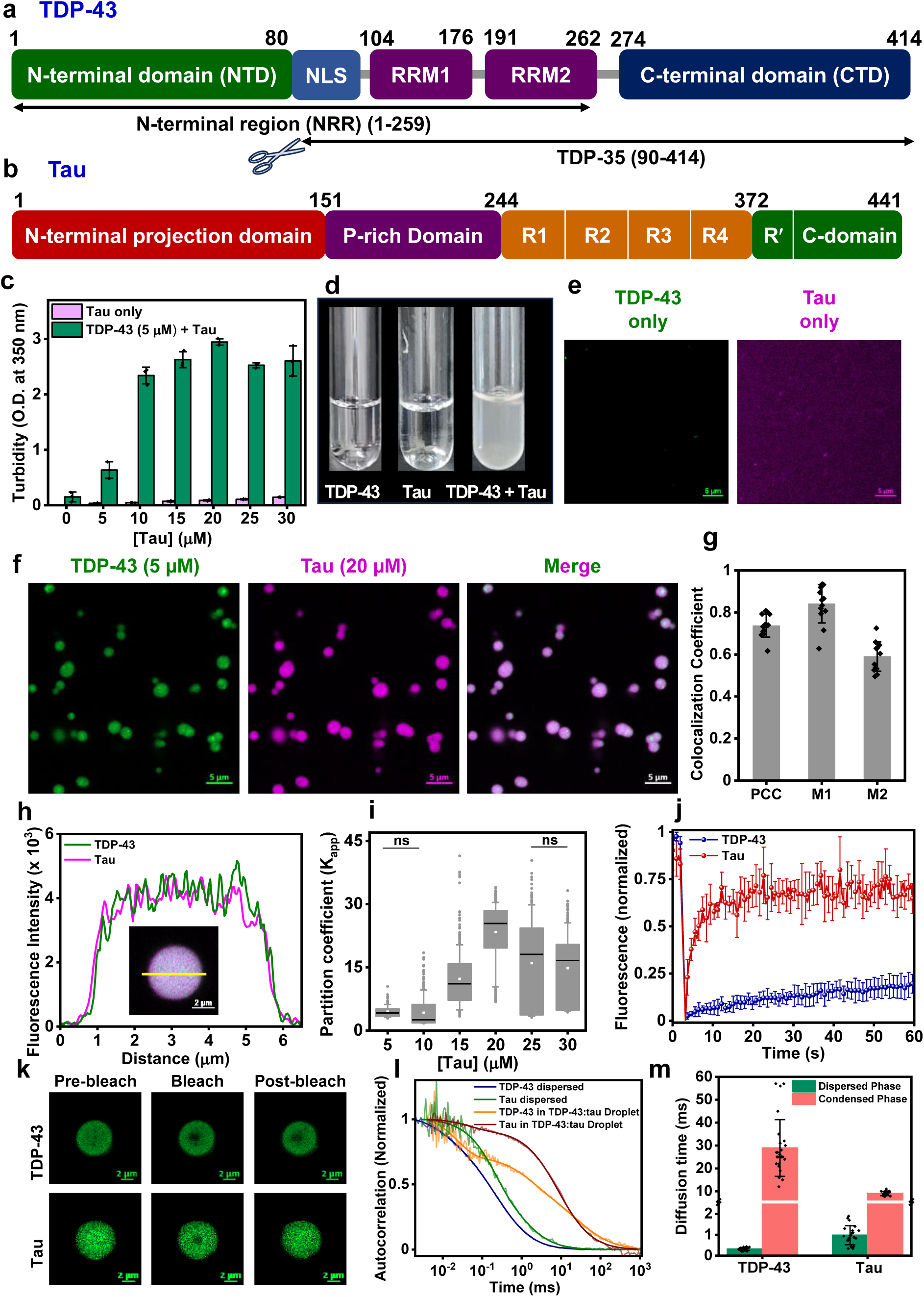
Heterotypic phase separation of TDP-43 and tau into heterogeneous condensates. Domain architecture of TDP-43 (**a**), highlighting the nuclear localization signal (NLS; residues 82-98) and the C-terminal TDP-35 fragment (90-414); and tau (**b**). **c.** The changes in turbidity as a function of tau concentration at a fixed TDP-43 concentration of 5 µM. Data represent mean ± SD (n = 3). **d.** Representative photographs for TDP-43 (5 µM), tau (20 µM), and TDP-43 + tau. Photographs were taken separately, then cropped and arranged. **e.** Representative two-color confocal Airyscan images showing no phase separation of TDP-43 (5 µM, AlexaFluor488) and tau (20 µM, AlexaFluor594). **f.** Phase separation upon mixing of TDP-43 and tau (20 mM HEPES, 15 mM NaCl, 1 mM DTT, pH 7.4). Scale bar = 5 µm. **g.** Colocalization plot showing the Pearson’s correlation and Manders’ colocalization coefficient for TDP-43:tau droplets. Data represent mean ± SD (n = 13). **h.** Line profile obtained for the fluorescence intensities of TDP-43 (green) and tau (magenta), showing colocalization within the heterotypic condensates (the droplet shown in the inset, scale bar = 2 µm). **i.** Apparent partition coefficient (K_app_) of TDP-43 within condensates, measured as a function of increasing tau concentrations. The data presented indicates the 25^th^ to 75^th^ percentile (box) and 10^th^ to 90^th^ percentile (whiskers). The median (line), mean (white square), and the outliers (gray dots) are also shown for n > 150 droplets for all concentrations, analyzed for more than seven independent samples. All data were statistically significant (p < 0.001) unless otherwise stated as ns. **j.** FRAP kinetics measured for AlexaFluor488-labeled TDP-43 (blue) and tau (red) within the co-phase-separated droplets. Data represent mean ± SD (n = 7). **k.** Representative images of AlexaFluor488-doped droplets during FRAP measurements (scale bar = 2 µm). **l.** Representative normalized FCS autocorrelation plots in dispersed and droplet phases. **m.** Mean diffusion times estimated from the FCS autocorrelation curves in the monomeric dispersed and droplet phases. Data represent mean ± SD (n = 22 and 25 independent repeats for the dispersed and droplet phases, respectively). See Table S2 for recovered parameters.

Under pathological conditions, hyperphosphorylation of tau causes its detachment from microtubules, resulting in the accumulation of the unbound tau within cytoplasmic assemblies, which eventually facilitates self-assembly into paired helical filaments (PHFs) and neurofibrillary tangles (NFTs)^44–46^. These deposits are identified as the neuropathological lesions implicated in AD and a range of other tauopathies. The interaction between TDP-43 and tau is known to compound the toxicity accompanied by early-onset and severe disease manifestations in AD patients. TDP-43 proteinopathy, which has been studied individually as the original hallmark of ALS and FTLD-TDP, has now emerged as a co-pathology in 50-70 % of AD patients^47–51^. TDP-43 pathology affecting older adults, termed as LATE-NC (limbic-predominant age-related TDP-43 encephalopathy neuropathological change), is now observed in association with AD neuropathological change, leading to more severe disease manifestations than those attributable to either pathology in isolation^52,53^. In particular, observations from recent neuropathological staging studies revealed a strong association between the co-deposition of hyperphosphorylated TDP-43 inclusions with tau NFTs and exacerbated pathological outcomes, including accelerated memory decline and greater hippocampal atrophy^54,55^. Here, using single-molecule FRET, in conjunction with other tools and cellular studies, we elucidate the molecular drivers of complex phase transitions mediated via domain-specific synergistic interactions between TDP-43 and tau. Our findings provide a generic mechanistic framework of abnormal heterotypic phase transitions associated with the progression of fatal neurodegenerative diseases.

## Results

### TDP-43 and tau undergo heterotypic phase separation into binary condensates

As a prelude, we first performed phase separation assays of recombinantly expressed and purified TDP-43 and tau using an *in vitro* cell-free reconstitution approach. Droplet formation was induced by cleaving the MBP tag from TDP-43 using TEV protease (1:10, TEV:TDP-43). We chose a buffer condition at physiological pH (20 mM HEPES, 15 mM NaCl, 1 mM DTT, pH 7.4), at which TDP-43 and tau do not undergo homotypic phase separation. Our turbidity measurements and confocal imaging confirmed that under these conditions, neither TDP-43 nor tau alone formed droplets (Fig. 1c-e). Next, we performed turbidity titration experiments using increasing concentrations of tau and by fixing the TDP-43 concentration at 5 µM. As the tau concentration increased, we observed a concentration-dependent increase in turbidity that saturated at a molar ratio of 1:3 (5 µM TDP-43 and 15 µM tau), indicating the formation of mesoscopic assemblies (Fig. 1d). To further confirm condensate formation, we performed two-color confocal Airyscan microscopy imaging. For visualization, we used AlexaFluor488-and AlexaFluor594-maleimide-labeled single-cysteine variants of TDP-43 and tau, respectively. Droplet reactions were doped with 1 % labeled protein, and both proteins showed clear enrichment within the droplet phase, indicating colocalization of TDP-43 and tau within these co-phase-separated condensates (Fig. 1f). Pearson’s correlation and Manders’ colocalization coefficients (Fig. 1g) and fluorescence intensity profiles (Fig. 1h) obtained for TDP-43 (5 µM) and tau (20 µM) further verified the strong spatial correlation between both proteins within these heterotypic condensates. Next, to probe the effect of varying protein stoichiometry, we aimed to determine the relative partitioning of TDP-43 within these condensates. Based on the relative fluorescence intensities measured in our two-color confocal imaging (Supplementary Fig. 1c), we estimated the apparent partition coefficient (Fig. 1i) for TDP-43 and observed an initial rise, followed by a gradual dip, indicating saturation of the heterotypic TDP-43:tau interactions at a molar ratio of 1:4, which was then fixed as the reaction stoichiometry for our subsequent experiments. The dip in the plot at higher stoichiometries could indicate the predominance of homotypic tau interactions over heterotypic TDP-43:tau contacts, corroborating the plateau in the concentration-dependent turbidity values (Fig. 1c).

Next, to probe the fluidity of these assemblies, we then set out to perform fluorescence recovery after photobleaching (FRAP) measurements using both AlexaFluor488-labeled TDP-43 and tau (Fig. 1j, k). Droplet reactions were separately doped with either labeled protein, and the fluorescence recovery of a submicron-sized region within the condensates was monitored post-bleaching. In the case of tau, FRAP measurements showed rapid, near-complete recovery, suggesting a highly dynamic, liquid-like nature of tau within these heterotypic condensates. In contrast, TDP-43 showed minimal recovery, indicating a predominantly immobile, gel-like behavior within the condensed phase, similar to that shown previously for homotypic TDP-43 assemblies^56^. This can be partially attributed to a dense intermolecular network of TDP-43 or to higher-order assembly formation, consistent with the oligomerization behavior of TDP-43^57,58^. FRAP measurements revealed the distinct differences in the mobility of TDP-43 and tau. We then employed fluorescence correlation spectroscopy (FCS) measurements to further quantify the diffusional properties of the polypeptide chains in these condensates. The diffusion times recovered from the FCS autocorrelation curves revealed fast diffusion for TDP-43 (∼ 0.3 ms) and slightly slower diffusion for tau (∼ 1 ms) in the monomeric dispersed phase (Fig. 1l, m, Table S2). Upon co-phase separation, both tau and TDP-43 experienced much slower diffusion within condensates. Tau exhibited a ∼ 9-fold slower diffusion time (∼ 9 ms), whereas, TDP-43 showed markedly slower diffusion (∼ 30 ms), indicating a ∼ 100-fold increase in the diffusion time within these heterotypic condensates (Fig. 1m). The diffusion times obtained within the droplet phase indicated a dynamic, liquid-like behavior for tau whereas TDP-43 existed in a significantly slower and less mobile state within the droplets. This highly reduced mobility, also consistent with FRAP measurements, suggests the formation of a dense network having long-lived intermolecular interactions within these heterotypic condensates. Taken together, this set of results demonstrates a concentration-dependent phase separation of TDP-43 and tau into heterotypic condensates with a protein-dense, viscoelastic interior. Since sequence-dependent interactions govern the emergent material properties of such condensates, we next set out to identify the molecular drivers of TDP-43:tau co-phase separation.

### Electrostatic and hydrophobic interactions drive co-phase separation of TDP-43 and tau

We next sought to discern the nature of the molecular interactions underlying this heterotypic co-condensation. We began by identifying the drivers underlying the multivalency and fuzzy interaction propensity of the two polypeptide chains. Our primary sequence analysis revealed a heterogeneous amino acid composition (Supplementary Fig. 2a) for both proteins with opposite charges at physiological pH. TDP-43 carries a net negative charge (-5), whereas tau is weakly positively charged (+ 2.5) under our experimental conditions. This complementary charge profile of these proteins hinted at a role of electrostatic interactions driving heterotypic phase separation. A close look at the charge patterning using the net-charge per residue (NCPR) analysis^59^ revealed a clustering of charged residues across the NTD and RRMs for TDP-43 (Supplementary Fig. 2b). In tau, on the other hand, the acidic residues are enriched in the N-terminal projection domain, whereas the proline-rich and MTBR domains contain blocks of basic residues. Based on these observations, we further aimed to determine the role of electrostatic interactions in promoting intermolecular contacts between TDP-43 and tau. We set up droplet reactions in the presence of increasing salt concentrations and observed a reduction in both the number and size of the phase-separated species, followed by complete dissolution of the condensates at a higher salt concentration (Fig. 2a and Supplementary Fig. 2c). We note that co-phase separation was also observed at the physiological salt concentration (150 mM NaCl) in the presence of a crowding agent (see later). These observations indicated that TDP-43:tau co-phase separation is mediated by electrostatic interactions that are weakened at higher salt concentrations due to increased charge screening.

**Figure 2:**
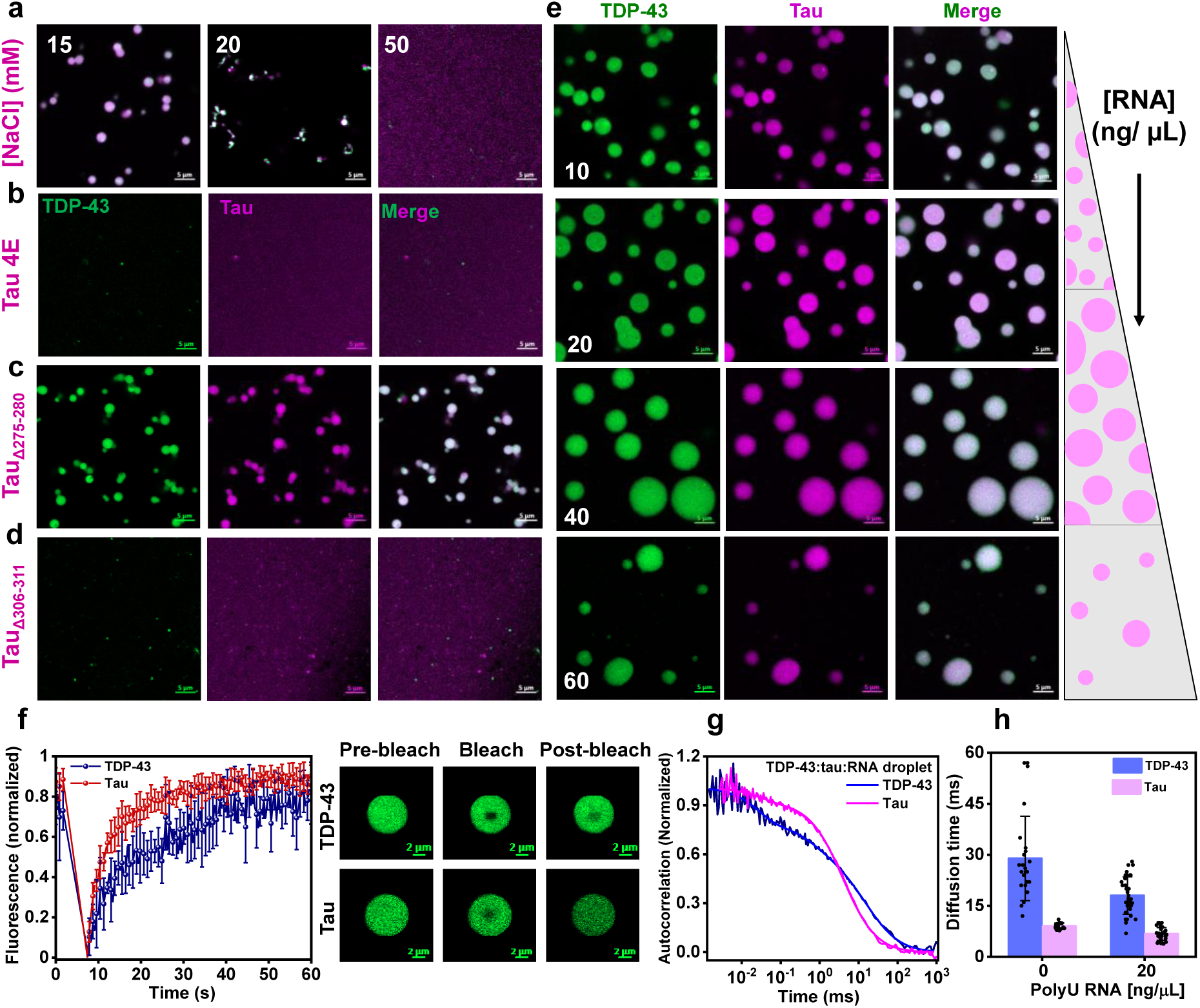
Interplay of electrostatic and hydrophobic interactions in TDP-43:tau co-condensation.a. Salt-dependent merged Airyscan confocal images of TDP-43 (5 µM) and tau (20 µM) showing dissolution of droplets at high concentration of NaCl. Representative two-color images of heterotypic condensation of TDP-43 (5 µM) with phosphomimetic tau (**b)**, tau_Δ275-280_ (20 µM) (**c**), and tau_Δ306-311_ (20 µM) (**d**), highlighting the role of electrostatic and hydrophobic contacts in driving TDP-43:tau co-condensation. **e.** RNA-driven tuning of the phase behavior of showed by the two-color Airyscan confocal imaging. Scale bar = 5 µm for all images (**a-e**). **f.** FRAP kinetics obtained for TDP-43 (blue) and tau (red) within condensates formed in the presence of 20 ng/µL PolyU RNA. Data represent mean ± SD (n = 5 and 8 independent repeats for TDP-43 and tau, respectively). Representative images from FRAP measurements are shown in the right panel (scale bar = 2 µm). Representative normalized FCS autocorrelation curves (**g**) and the corresponding diffusion times (**h**) recovered from the FCS curves for AlexaFluor488-labeled TDP-43 and tau inside condensates formed with 20 ng/µL PolyU RNA. Data represent mean ± SD (n = 34 and 38 independent repeats for TDP-43 and tau, respectively). Data for 0 ng/µL are the same as shown in Fig. 1m and are shown here for comparison. See Table S2 for recovered parameters.

Additionally, changes in protein net-charge due to posttranslational modifications, such as phosphorylation, can modulate protein phase behavior. For instance, tau phosphorylation is a well-known hallmark and a potent driver of the aberrant phase transition, a mechanism underlying cellular neuropathology^44–46^. This further motivated us to investigate the effect of tau phosphorylation on these heterotypic condensates. To this end, we employed a phosphomimetic tau mutant (Tau 4E), which recapitulates phosphorylation by microtubule affinity-regulating kinases in cells and harbors four mutations that introduce glutamate residues. Similar to the effects observed upon increasing ionic strength, incorporation of these additional negative charges into the tau polypeptide chain resulted in complete dissolution of the condensates (Fig. 2b), further supporting the role of electrostatic interactions in driving TDP-43:tau co-phase separation. Furthermore, both TDP-43 and tau comprise segments of hydrophobic residues (Supplementary Fig. 2a), underlining the potential of hydrophobic interactions as molecular drivers. To test this, we disrupted weak hydrophobic contacts using 1,6-hexanediol (1,6-HD). Our turbidity measurements and two-color confocal Airyscan imaging revealed a reduction in overall phase separation propensity, resulting in the formation of smaller assemblies (Supplementary Fig. 2d,e). However, 1,6-HD failed to completely abolish TDP-43:tau co-condensation even at higher concentrations (10 %), indicating that in the absence of hydrophobic interactions, electrostatic interactions are capable of forming smaller assemblies. To delve deeper into the contribution of hydrophobic interactions, we utilized two tau mutants with deletions of either of the hydrophobic hexapeptide repeats namely, Δ275-280 (termed PHF6*) and Δ306-311 (termed PHF6) that are located in the MTBR domain (residues 244-372) and are associated with homotypic tau phase separation and amyloid formation. Deletion of the first repeat (Δ275-280) region did not alter the heterotypic phase separation with TDP-43 (Fig. 2c). Surprisingly, deletion of the second hexapeptide repeat (Δ306-311), with the first repeat intact (275-280), resulted in complete abrogation of phase separation with TDP-43, as observed in our two-color Airyscan confocal imaging (Fig. 2d). This observation is in line with the indispensable role of this region in driving tau condensation, suggesting a role of tau self-interaction in driving heterotypic TDP-43:tau condensation^26^. These results suggest that both electrostatic and hydrophobic interactions are necessary for the TDP-43:tau co-phase separation.

### RNA tunes the heterotypic phase behavior of TDP-43 and tau

TDP-43 is an RNA-binding protein containing RRMs and is involved in mRNA regulation, processing, transport, as well as transcription and translation. The phase-separating fate of such an RNA-binding protein is also determined by RNA^12,60–62^. Mutations that result in RNA-binding-deficient mutants disrupt the balance between the nuclear and cytoplasmic protein pools, invariably leading to pathological conditions via the loss of function in the nucleus or the gain of function in the cytoplasm. Additionally, both TDP-43 and tau are associated with RNA-rich cytoplasmic stress granules, and thus, RNA emerges as a potential regulator of their co-phase separation^36–38,63^. In this context, we aimed to decipher the role of RNA in regulating the co-condensation of TDP-43 and tau. We first measured turbidity as a function of increasing concentration of polyU RNA. At lower RNA concentrations, we observed a slight increase in turbidity, followed by a plateau at 40 ng/µL RNA, after which phase separation decreased and was abolished at 100 ng/µL RNA (Fig. S2f). The initial rise followed by a gradual inhibition of droplet formation can be explained by successive charge screening and inversion, which recapitulate the electrostatically driven RNA-induced reentrant phase behavior observed for other RNA-binding proteins^62^. Confocal imaging confirmed the formation of liquid-like ternary condensates, with RNA promoting the formation of larger droplets at low concentrations and dissolution at higher RNA concentrations (Fig. 2e). Next, to probe the interior of TDP-43:tau:RNA ternary condensates, we performed FRAP measurements and monitored the recovery of both the proteins within condensates formed at lower RNA concentration (20 ng/µL) (Fig. 2f). Sub-stoichiometric concentrations of RNA that promote phase separation can facilitate dense protein-RNA electrostatic crosslinking characterized by long-lived intermolecular contacts. Such enhanced physical crosslinking can increase the viscoelasticity and restricted molecular mobility within condensates^60,62,64^. In contrast, our FRAP and FCS results demonstrated that TDP-43 exhibited highly dynamic behavior within these TDP-43:tau:RNA ternary condensates (Fig. 2f-h). A more efficient FRAP recovery and faster diffusion time of TDP-43 indicated that RNA can alter protein-protein interactions and buffer TDP-43 phase separation by modulating both homotypic and heterotypic interactions. Taken together, these findings indicated RNA-dependent tuning of protein-protein interactions that modulate TDP-43:tau co-condensation. Such a regulation of condensate properties suggests a possible role of RNA in rewiring the intermolecular protein network within RNA-rich membraneless compartments. Next, we aimed to discern the role of domain-specific interactions between TDP-43 and tau in their co-phase separation.

### The N-terminal domain and RRMs of TDP-43 are crucial for co-condensation

Both TDP-43 and tau exhibit a multi-domain architecture (Fig. 1a,b), with widely different sequence attributes and functions assigned to each domain. Hence, we next probed the domain-specific facilitators of the intermolecular interaction network driving complex two-component phase separation. To this end, we utilized two truncation variants of TDP-43, one expressing the N-terminal folded domain and the two RRMs (residues 1-259, termed NRR) and another the disordered C-terminal domain (residues 267-414, termed CTD) (Fig. 1a). The NRR comprises the folded domain along with the two RRMs, which are known to mediate dimerization and RNA binding, respectively, whereas the CTD represents a low-complexity, prion-like domain responsible for phase separation/aggregation, and harbors disease-associated mutations of TDP-43^36,65^. We began by investigating the ability of either domain to drive co-condensation with tau independently. Strikingly, the NRR alone was sufficient to drive robust phase separation with tau, forming condensates comparable to those of full-length TDP-43 (Fig. 3a, top panel). In contrast, the isolated CTD had a reduced propensity to undergo co-phase separation with tau compared to full-length TDP-43, evident by smaller droplets with the CTD (Fig. 3a, bottom panel). These observations indicate that, in the context of heterotypic co-condensation with tau, the chief determinants of phase separation reside within the folded NTD and the two RRMs (collectively termed NRR) of TDP-43. However, we note here that the CTD of TDP-43 alone undergoes homotypic phase separation and can recruit tau into these condensates. This observation is consistent with a previous study performed in the presence of crowding agents^66^.

**Figure 3:**
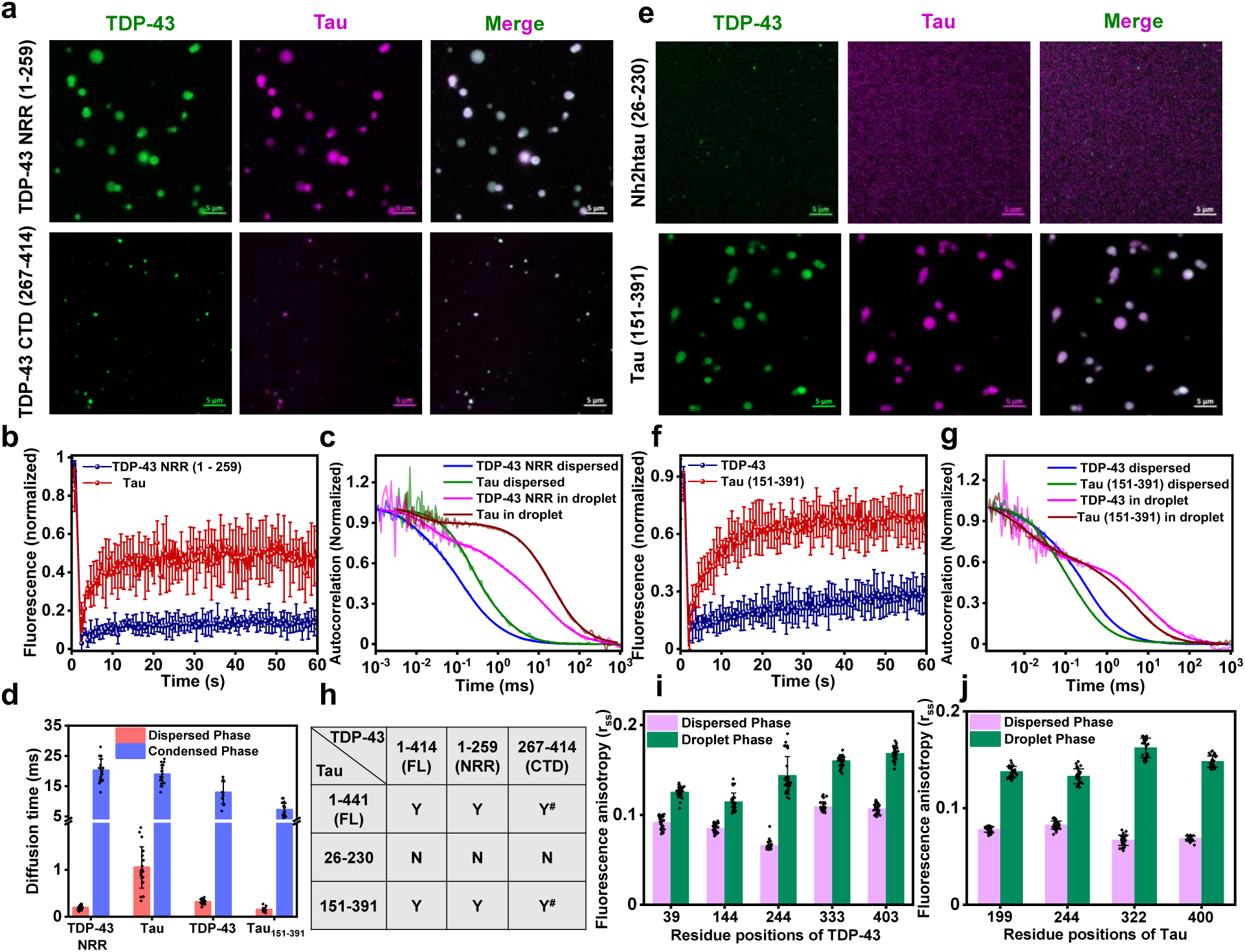
Domain-specific interactions underlying TDP-43:tau co-condensation. **a.** Representative two-color confocal Airyscan images of droplet reactions upon mixing tau (20 µM) with TDP-43 NRR (5 µM, top panel) and TDP-43 CTD (10 µM, bottom panel) under our reaction conditions. For visualization, reactions were doped with 1 % AlexaFluor488-labeled TDP-43 variants and 1 % AlexaFluor594-labeled full-length tau. Scale bar = 5 µm. **b.** FRAP kinetics averaged over multiple droplets formed with TDP-43 NRR and tau. Data represent mean ± SD (n = 6 independent repeats for both TDP-43 NRR and tau). **c.** Normalized autocorrelation curves obtained in the dispersed and droplet phases. **d.** Diffusion times estimated from the FCS autocorrelation curves in the dispersed and condensed phases. Data represent mean ± SD (n = 20 and 19 independent repeats for TDP-43 NRR and tau, respectively, in the dispersed phase; n = 18 and 20 independent repeats for TDP-43 NRR and tau, respectively, in the droplet phase). For tau_151-391_ and TDP-43, n = 18 in the dispersed phase and n = 10 and 16 for TDP-43 and tau_151-391_, respectively, in the droplet phase. See Table S2 for recovered parameters. **e.** Representative two-color confocal Airyscan images of phase separation of TDP-43 (5 µM) with Nh2htau (tau_26-230_) (20 µM, top panel) and tau_151-391_ (20 µM, bottom panel). Scale bar = 5 µm. **f.** FRAP kinetics measured for AlexaFluor488-labeled TDP-43 and tau_151-391_ within their co-phase-separated droplets. Data represent mean ± SD (n = 9 and 10 independent repeats for TDP-43 and tau_151-391_, respectively. **g.** Representative normalized FCS autocorrelation plots obtained in the dispersed and droplet phases. **h.** A tabular summary of domain-specific interactions necessary for driving TDP-43:tau co-condensation (FL: full-length): Y (yes) and N (no) indicate whether co-phase separation was observed or not (^#^ small droplets formed via homotypic phase separation of TDP-43 CTD recruit tau). Steady-state fluorescence anisotropy measured for various F5M-labeled single-cysteine variants of TDP-43 (**i**) and tau (**j**) in the dispersed phase and within single droplets of TDP-43:tau. Data represent mean ± SD (n = 30 independent repeats for both dispersed and droplet phases).

Next, to probe the material properties of these NRR-driven condensates, we performed FRAP measurements within the condensates and observed a slow, limited recovery (∼ 15 %) for TDP-43 NRR, similar to the behavior of full-length TDP-43 (Fig. 3b). However, tau recovery decreased dramatically to ∼ 50 %, compared with the near-complete, rapid recovery observed in droplets formed with full-length TDP-43 (Fig. 3b). Our FCS measurements further demonstrated a nearly 100-fold increase in the diffusion times of TDP-43 NRR (from ∼ 0.2 ms to ∼ 20 ms) and a ∼ 19-fold increase in tau (from ∼ 1 ms to ∼ 19 ms) from the dispersed phase to the condensed phase (Fig. 3c,d). Interestingly, tau diffusion was ∼ 2 times slower compared to full-length TDP-43:tau condensates, suggesting a denser, less dynamic network. Furthermore, previous studies showed that the disordered CTD facilitates transient α-helix-mediated TDP-43 self-association and, in turn, head-to-tail oligomerization^65^. Therefore, we speculate that the absence of self-association-driving CTD interactions preferentially drives TDP-43 NRR towards heterotypic contacts with tau, as reflected in the slower diffusion of tau within these condensates. To further probe the role of domain-specific interactions in TDP-43, we employed three different point mutations of TDP-43: a phosphomimetic mutant (S48E) in the NTD and two disease-associated point mutations (Q331K and G335D) known to alter the helix-forming ability of TDP-43 CTD^58,65^. The S48E substitution is known to disrupt the β-sheet interface that mediates TDP-43 dimerization, thereby reducing overall self-association. Despite the additional negative charge and reduced propensity for homotypic contacts, the phosphomimetic mutant showed significantly reduced phase separation with the positively charged tau (Supplementary Fig. 3a,b). This observation was contrary to the enhanced heterotypic interactions observed upon removal of the self-associating CTD, indicating the specificity of the NTD-driven contacts beyond electrostatics. The mutations in the CTD (Q331K and G335D) did not significantly alter the affinity for tau, as seen in our turbidity and imaging experiments (Supplementary Fig. 3a,c,d). These findings indicated that the NRR of TDP-43 is primarily responsible for TDP-43:tau co-condensation.

### The central region of tau potentiates TDP-43:tau co-condensation

Next, to determine the interacting domains within tau, we employed two pathologically relevant truncation mutants of tau: a mutant expressing part of the N-terminal projection domain and the central proline-rich region (Nh2htau, residues 26-230) and another variant comprising the proline-rich domain and the C-terminal repeat regions (tau_151-391_ residues 151-391) (Fig. 1a). The N-terminal domain is highly disordered, whereas the relatively conformationally-rigid proline-rich motif can facilitate interactions with diverse binding partners and harbors multiple phosphorylation sites that allow dynamic regulation of tau phase separation. The negatively charged N-terminal region and the proline-rich region alone failed to undergo phase separation with TDP-43 under our reaction conditions (Fig. 3f, top panel). As expected for highly positively charged tau_151-391_ (net-charge ≍ + 24), it exhibited droplet formation with negatively charged TDP-43 (Fig. 3f, bottom panel). Our FRAP measurements revealed a slight improvement in TDP-43 recovery with tau_151-391_ compared with TDP-43:tau droplets (Fig. 3g). FCS measurements revealed a faster diffusion for tau_151-391_ compared to full-length tau in both the dispersed (∼ 0.15 ms) and the condensed (∼ 7 ms) phases (Fig. 3d,g and Table S2). Interestingly, TDP-43 diffusion was more than two times faster in these droplets (∼ 13 ms) than that in the droplets with full-length tau (∼ 30 ms), suggesting a relatively dynamic droplet interior. This can be attributed to the absence of the negatively charged N-terminal projection domain of tau, which presumably participates in electrostatic interactions with the positively charged RRM regions of TDP-43, contributing to a dense physical cross-linking within the condensates. Our co-phase separation experiments on individual domains of both TDP-43 and tau revealed the role of each of these domains in driving condensate formation (Fig. 3h). We also probed the domain-specific conformational flexibility of the polypeptide chains by measuring single-droplet fluorescence anisotropy that captures the extent of rotational dynamics in a site-specific manner. An increase in the fluorescence anisotropy of fluorescein-5-maleimide (F5M)-labeled at various single-cysteine positions across different domains of both proteins was observed in the droplet phase, indicating a restricted rotational mobility in the protein-rich condensed phase (Fig. 3i,j and Supplementary Fig. 4a,b). Our anisotropy results showed that RRM2 of TDP-43 (around residue 244) and the MTBR region of tau (around residue 322) exhibited the most significant rotational restriction in the condensed phase, underscoring the vital roles of these regions in promoting co-phase separation. Together, this set of findings characterize the domain-specific interactions which, upon disease-associated truncations, can alter condensate physical properties, potentially predisposing these assemblies to convert into pathological co-aggregates.

### Single-molecule FRET reveals structural unwinding of TDP-43 during co-condensation

Next, we asked what critical molecular events drive TDP-43:tau co-condensation. To this end, we employed single-molecule FRET (Förster resonance energy transfer) that offers a powerful tool to directly discern structural subpopulations and characterize the changes in conformational distributions upon phase separation^26,67,68^. We aimed to study the structural changes of TDP-43 that facilitate its co-phase separation with tau. Our results described in the previous sections established that the N-terminal region (NRR; residues 1-259) provides the primary electrostatic contacts that drive TDP-43:tau co-condensation. Thus, we set out to perform single-molecule FRET to characterize the conformational changes in the NRR, comprising the NTD and the two RRMs, during the co-phase separation of TDP-43 and tau.

To achieve this, we first prepared a null-cysteine mutant of TDP-43 that retained the overall secondary structural elements, as evidenced by our circular dichroism (CD) measurements (Supplementary Fig. 5a). This null-cysteine mutant was further used to create two dual-cysteine variants spanning the NTD and the RRM1 (39C-144C, a separation of 105 residues) and encompassing the RRM1 and RRM2 (144C-244C, a separation of 100 residues) (Fig. 4a). These constructs were labeled using thiol-active donor-acceptor FRET pairs, namely AlexaFluor488 maleimide and AlexaFluor594 maleimide, respectively. We then performed single-molecule FRET experiments (100-200 pM, 20 mM HEPES, pH 7.4) for freely diffusing fluorescently labeled TDP-43 molecules in the dispersed phase using a two-color pulsed-interleaved excitation (PIE) mode on a time-resolved confocal microscope, as described by us previously^26,68^. Such low concentrations used in our single-molecule experiments permit us to record the donor and acceptor fluorescence bursts from the monomeric form of the protein diffusing through the tiny confocal volume. Single-molecule PIE-FRET histogram for the 39-144 construct exhibited a bimodal distribution with a dominant FRET efficiency peak ∼ 0.8 with a smaller subpopulation at ∼ 0.6 (Fig. 4b). Since the wild-type TDP-43 can potentially exist in a monomer-dimer equilibrium under physiological conditions^58^, we introduced phosphomimetic substitutions (S333D and S342D) shown to stabilize the monomeric state^69^ to minimize any contribution of intermolecular FRET from higher-order species. The FRET histogram of this 39C-144C construct harboring S333D and S342D mutations exhibited a near-identical bimodal FRET distribution (Supplementary Fig. 5b), indicating that the two FRET efficiency peaks indeed arose from monomeric structural subpopulations. These structural states can potentially undergo conformation exchange on a much slower time scale than the diffusion timescale. We next set out to determine the changes in the conformational distribution of TDP-43 upon co-condensation with tau.

**Figure 4:**
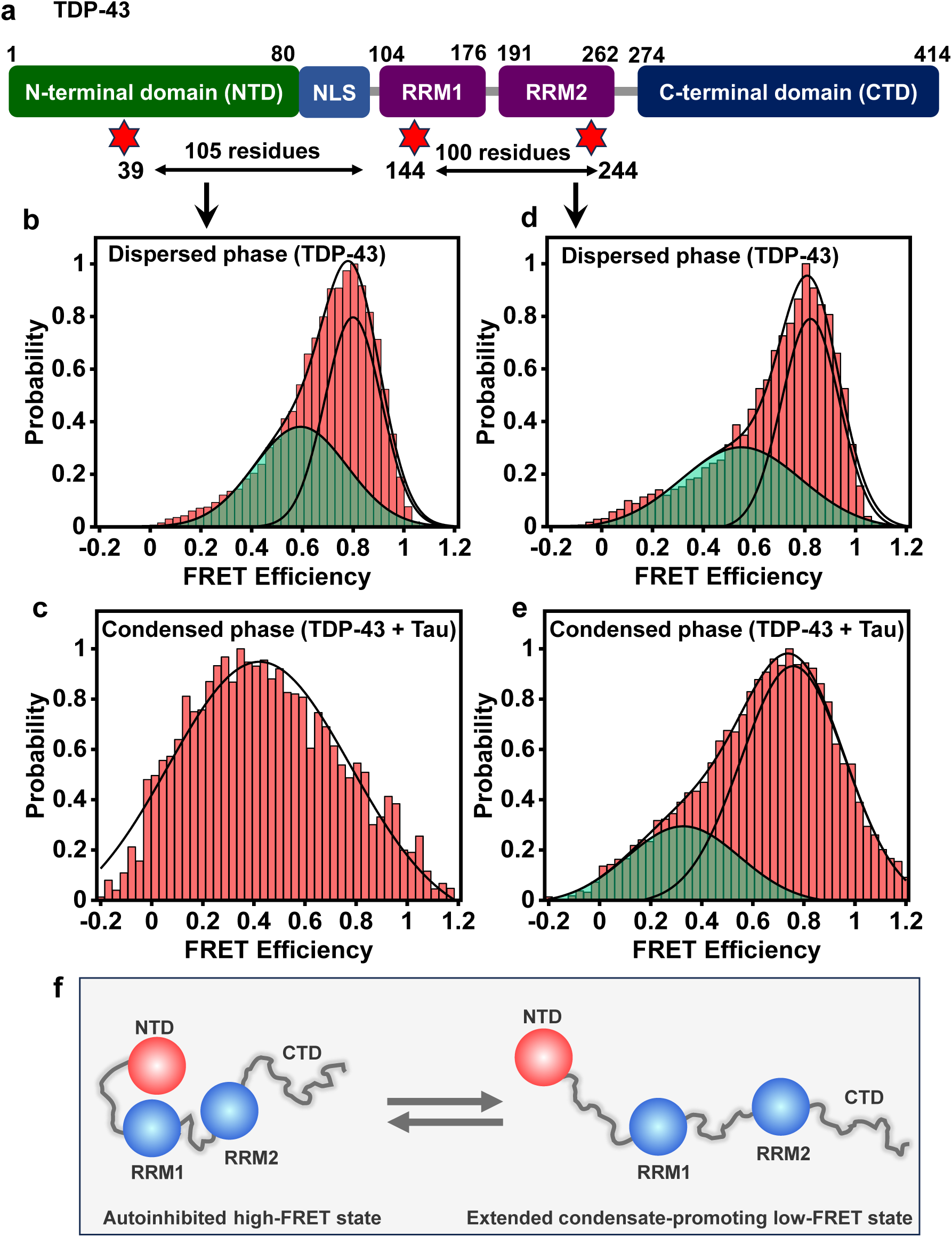
Single-molecule FRET reveals conformational switch accompanying TDP-43:tau co-condensation. **a.** TDP-43 single-molecule FRET construct details highlighting the sites involved in FRET. Single-molecule FRET (donor: AlexaFluor488; acceptor: AlexaFluor595) histograms with fits obtained for the 39-144 construct in the monomeric dispersed phase of TDP-43 (**b**) and droplet phase of TDP-43 and tau (**c**). Single-molecule FRET histograms with fits obtained for the 144-244 construct in the monomeric dispersed phase of TDP-43 (**d**) and droplet phase of TDP-43 and tau (**e**). The total number of events obtained for all single-molecule FRET measurements was always > 20,000. **f**. A schematic of structural subpopulations based on single-molecule FRET results showing an equilibrium of compact autoinhibited high-FRET state and extended condensate promoting low-FRET state that can participate in homotypic and heterotypic multivalent interactions leading to phase separation.

For single-molecule FRET measurements in the condensed phase, we focused the lasers inside large immobilized droplets (∼ 2 µm from the surface into the droplet), as previously described by us^26,68^. For these experiments, the concentrations of TDP-43 and tau were 5 µM and 20 µM, respectively, while the dual-labeled TDP-43 concentration was maintained at ∼ 30 pM. Upon co-phase separation with tau, the 39C-144C construct of TDP-43 exhibited an overall shift towards a broad, lower, unimodal FRET efficiency distribution centered around ∼ 0.4, indicating structural unwinding of a part of the NRR comprising the NTD and RRM1 (Fig. 4c). Similarly, the region spanning RRM1 and RRM2 (FRET pair 144-244) also exhibited two subpopulations and underwent structural unwinding, as shown by the slightly lower shift and broadening of the FRET efficiency peaks (Fig. 4d,e). We, however, would like to note that the changes in the peak width can be attributed to a combination of the photon shot noise and inherent conformational heterogeneity. Taken together, our single-molecule FRET results indicated that the central part of TDP-43, comprising RRM1 and RRM2, unraveled during co-phase separation with tau, albeit to a lesser extent than the segment consisting of the NTD and RRM1. Our findings suggested the coexistence of two distinct conformational subpopulations. The dominant high-FRET state represents a compact autoinhibitory state of NRR in which the acidic NTD can participate in intramolecular electrostatic interactions with RRM1 and RRM2, precluding TDP-43 from undergoing phase separation (Fig. 4f). The extended low-FRET state with a lesser extent of intramolecular interactions can engender multivalent intermolecular contacts leading to phase separation. Tau alters the balance between intra-and intermolecular interactions, creating a preference for extended condensate-promoting conformers of TDP-43 for heterotypic phase separation. Such a conformational expansion coupled with an increase in structural heterogeneity can unravel the compact, self-interacting, monomeric form of TDP-43, allowing the formation of intermolecular physical crosslinks via electrostatic and hydrophobic interactions, ensuing heterotypic phase separation of TDP-43 and tau. We next asked if these TDP-43:tau co-condensates can undergo aberrant phase transitions into amyloid-like aggregates that are the hallmarks of pathological conditions.

### Liquid-to-solid phase transitions lead to the formation of amyloid-like cytotoxic species

To investigate whether TDP-43:tau heterotypic interactions persist under physiologically relevant conditions, we began by reconstituting phase separation in the presence of a molecular crowding agent, namely, polyethylene glycol (PEG 8000) and physiological salt concentration (150 mM NaCl). Upon mixing TDP-43 and tau in 10 % PEG and 150 mM NaCl, we observed a sharp increase in turbidity, indicating increased phase separation (Supplementary Fig. 6a). The formation of heterotypic droplets was confirmed by our two-color Airyscan confocal imaging, which showed colocalization of both TDP-43 and tau within these condensates (Fig. 5a). FRAP measurements within these droplets recapitulated the behavior of both proteins inside droplets formed in the absence of crowding agents and salt (Supplementary Fig. 6b). To verify the electrostatic nature of the molecular drivers underlying this co-condensation, we measured turbidity in droplet reactions as a function of salt concentration (Supplementary Fig. 6c). Similar to droplets formed under low-salt conditions, phase separation under physiological conditions also showed salt-dependent modulations, reiterating the electrostatic nature of these heterotypic assemblies. As shown previously, these weak, transient, multivalent interactions are responsible for maintaining the dynamism and reversibility of condensates that can gradually undergo an aberrant liquid-to-solid phase transition into aggregates^27–35^. We next set out to investigate such a phase separation-mediated aggregation of TDP-43:tau co-condensates.

**Figure 5:**
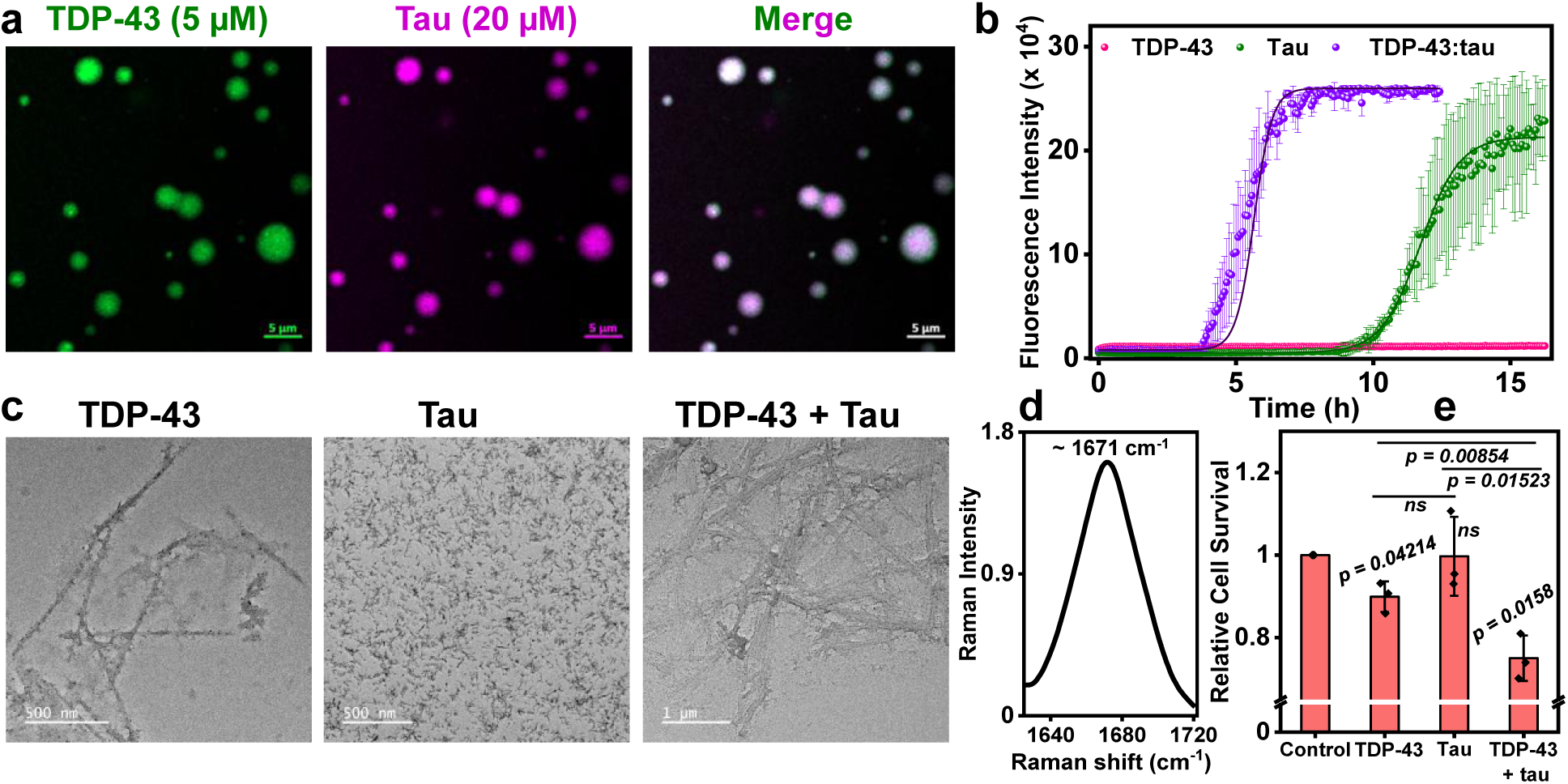
Phase separation-mediated transition to amyloid fibrils. **a.** Representative two-color confocal Airyscan image showing co-localization of TDP-43 and tau within heterotypic condensates under the physiological condition in the presence of a crowding agent (20 mM HEPES, 150 mM NaCl, 10 % PEG, pH 7.4). Scale bar = 5 µm. **b.** The ThT kinetics of droplets formed by TDP-43 (pink), tau (olive), and TDP-43:tau (purple) upon aging under stirring conditions. **c.** TEM images of the aggregated species formed by TDP-43 and tau after 16 h, and TDP-43:tau after 10 h. Scale bar = 500 nm for TDP-43 and tau alone condensates and 1 µm for TDP-43:tau condensates. **d.** Raman Amide I band obtained for phase separation-mediated TDP-43:tau co-aggregates obtained at the end point (10 h). **e.** Relative cell survival of HeLa cells 72 h after incubation with *in vitro*-formed aggregates, normalized to the untreated control. Data represent mean ± SD (n = 3 independent biological replicates). The p-values were determined using a paired t-test and are reported in the plots.

Homotypic and heterotypic phase separation of TDP-43 (5 μM) and tau (20 μM) was induced in the presence of PEG 8000 (20 mM HEPES, 150 mM NaCl, 10 % PEG, 1 mM DTT, pH 7.4) (Fig. 5a and Supplementary Fig. 6a), and the reactions were incubated under shaking conditions to facilitate the aging and liquid-to-solid transition of the assemblies. TDP-43 and tau are both known to form amyloid-like aggregates. Thus, to study the aggregation kinetics, we monitored the fluorescence of the amyloid-specific dye thioflavin T (ThT) (Fig. 5b). For homotypic TDP-43 condensates, we did not observe a significant increase in ThT fluorescence. This observation is consistent with previous studies showing that the amyloid aggregates formed by TDP-43 are not ThT-positive, despite harboring β-sheet-rich structures, due to the shielding of the amyloid core by a fuzzy coat comprising the N-terminal regions^70^. For tau-only droplets, the ThT fluorescence increased after a long lag time of ∼ 9 h, whereas in the case of co-phase-separated TDP-43:tau droplets, the lag time was considerably reduced to ∼ 4 h, indicating that TDP-43:tau co-condensates undergo facile conversion into ThT-positive aggregated species. To visualize these species, we performed transmission electron microscopy (TEM) on aggregates obtained at 10 h and 16 h in the heterotypic (TDP-43:tau) and homotypic (TDP-43 and tau alone) conditions, respectively. The TDP-43-only droplets matured into long, rod-like fibrillar structures, whereas the comparatively slower kinetics and lower ThT intensity originated from small, thin, protofibrillar structures formed in tau-only condensates (Fig. 5c). On the other hand, the aging of co-phase-separated droplets led to the formation of aggregates with properties intermediate between those formed by either protein individually, indicating the formation of co-aggregates mediated by co-condensation. TEM imaging revealed the formation of thick, dense, and bundled fibril-like structures morphologically closer to TDP-43 aggregates with ThT binding, similar to that of tau-only aggregates (Fig. 5b,c). Furthermore, we performed vibrational Raman spectroscopy to obtain structural insights into fibrillar architecture. Raman spectra revealed typical vibrational modes with prominent marker bands, including Amide I (1600-1700 cm^-1^) and Amide III (1250-1350 cm^-1^), which are attributed to the C=O stretching and a combination of C-N stretching and in-plane N-H bending modes of the polypeptide chain, respectively (Supplementary Fig. 6d)^71,72^. Both Amide I and Amide III indicated the presence of β-sheet-rich structure in aggregates. A characteristic Amide I band peak at ∼ 1670 cm^-1^, arising from the in-phase C=O stretching, indicated a typical cross-β architecture in the aggregates (Fig. 5d and Supplementary Fig. 6e-g, olive). Interestingly, a minor peak from the disordered conformation exhibited varying contributions, indicating the formation of structurally distinct fibrils upon TDP-43:tau co-aggregation (Supplementary Fig. 6e-g, red). Additionally, we determined the ratio of the tyrosine Fermi doublet (I_850_/I_830_) that increased from ∼ 0.7-0.8 (TDP-43-only and tau-only aggregates) to ∼ 1.5 (TDP-43:tau co-aggregates), indicating enhanced hydrogen bonding via tyrosine hydroxyl group in TDP-43:tau co-aggregates. Collectively, these results suggest that heterotypic condensates of TDP-43 and tau undergo maturation into amyloid-like co-aggregates that are distinct from the aggregates derived from individual proteins. Next, to test whether these co-aggregates could drive cellular toxicity, we monitored cell viability upon treatment with aggregates containing only TDP-43, only tau, or both TDP-43 and tau using a colorimetric MTT (3-(4,5-dimethylthiazol-2-yl)-2,5-diphenyltetrazolium bromide) assay. The relative survival of cells treated with the different aggregates was compared with that of untreated cells, indicating that TDP-43:tau co-aggregates elicited greater cytotoxicity than aggregates of individual proteins (Fig. 5e). Taken together, our findings suggested that heterotypic phase separation of TDP-43 and tau promotes the formation of structurally distinct, cytotoxic, amyloid-like co-aggregates that can potentially be important in the context of tau-associated neuropathological conditions. We next asked whether similar interactions could be observed within the cellular environment, particularly within RNA-rich stress granules that are enriched in both TDP-43 and tau.

### TDP-35 and tau are recruited into cytoplasmic granules during oxidative stress

TDP-43 is involved in several RNA-associated functions within the nucleus and is also associated with cytoplasmic stress granules, which recruit translationally halted mRNAs along with several other aggregation-prone, low-complexity domain-containing RBPs^36–38^. Furthermore, in some cases, these granules can also provide a hotspot for abnormal protein aggregation, leading to toxic species that mature into irreversible pathological deposits from postmortem ALS/FTLD specimens^29,30,73^. Additionally, tau has been shown to be associated with the formation and stabilization of stress granules^63^. Thus, we asked whether these stress granules could facilitate the co-assembly and phase transition of TDP-43 and tau inside cells. To this end, we began by transiently expressing N-terminally AcGFP-tagged TDP-43 and C-terminally mCherry-tagged tau in HeLa cells. To recapitulate the oxidative stress-induced cellular response and initiate stress granule formation, the cells were treated with 300 µM sodium arsenite for 1 h (Supplementary Fig. 7a,b). Due to the presence of bipartite NLS in the N-terminal part, TDP-43 exhibited a complete nuclear localization, whereas tau was distributed in the cytoplasm, as revealed by our two-color confocal imaging of cells co-expressing TDP-43 and tau (Fig. 6a). Recruitment into stress granules is shown to be dependent on the relocalization of TDP-43 from the nucleus to the cytoplasm either by employing a combination of stresses and pathological mutations^36–38^. Thus, in order to facilitate cytoplasmic localization of TDP-43 and to further recapitulate the cytoplasmic mislocalization preceding pathology, we chose to use TDP-35, which is a naturally occurring, ∼ 35 kDa C-terminal fragment (residues 90-414. Fig. 1a), generated as a result of caspase-mediated cleavage in the N-terminal domain of TDP-43^74,75^ and is highly enriched in specific brain regions of patients with ALS/FTLD^39,43^. We, therefore, carried out the transient expression of this pathologically relevant construct of TDP-43 (TDP-35) along with tau.

**Figure 6:**
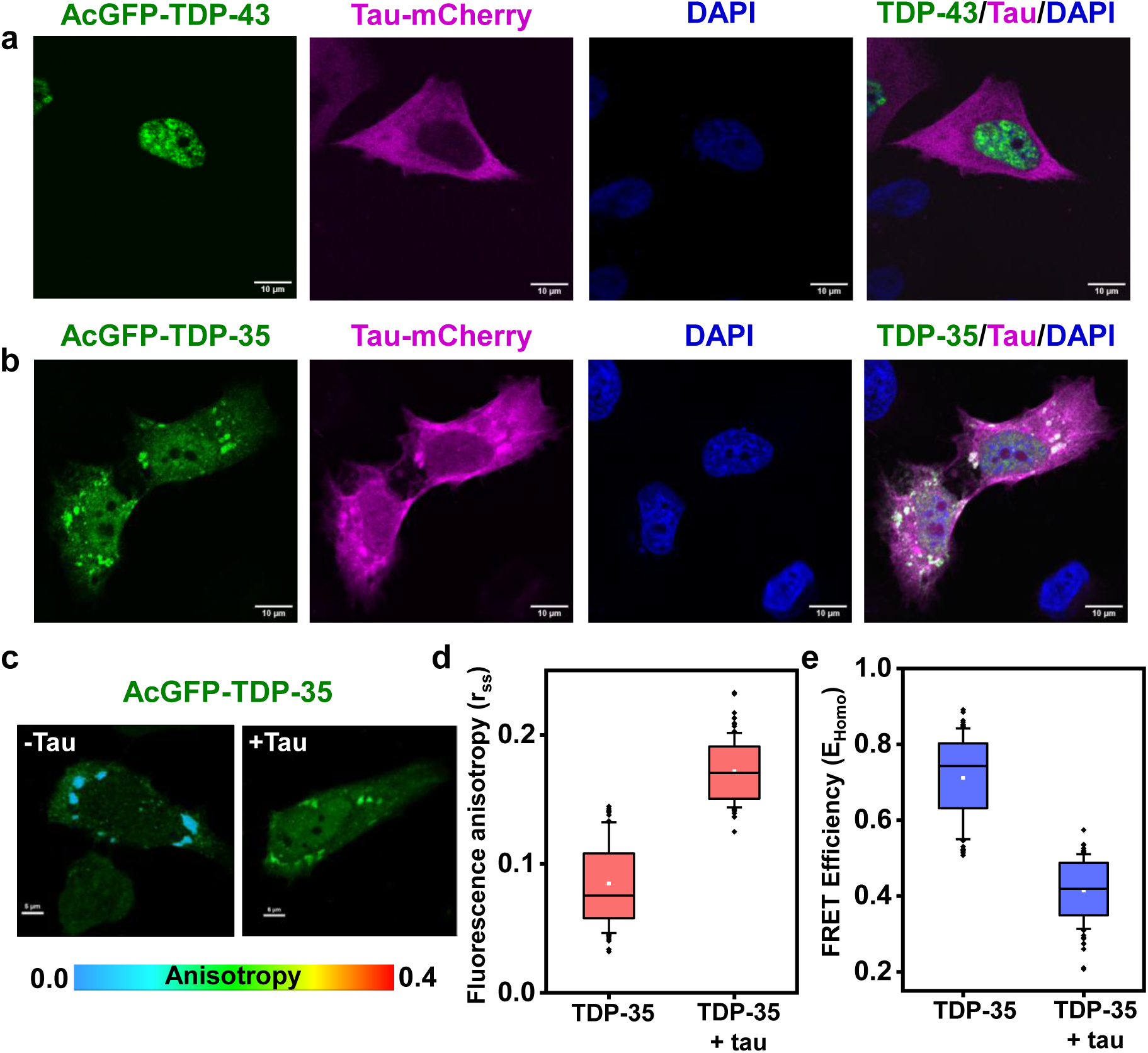
Association of tau with pathological TDP-35 within stress-induced cytoplasmic granules. **a.** Representative two-color confocal images of HeLa cells transiently overexpressing AcGFP-TDP-43 (green) and tau-mCherry (magenta) with nuclear (DAPI) staining and treated with 300 µM sodium arsenite to recapitulate cellular oxidative stress. TDP-43 localizes to the nucleus, whereas tau remains diffused in the cytoplasm. **b.** Representative two-color confocal image of HeLa cells overexpressing AcGFP-TDP-35, a pathological variant of TDP-43, along with tau-mCherry. Upon oxidative stress conditions, TDP-35 forms cytoplasmic granules within which tau-mCherry also colocalizes. Scale bar = 10 µm for all images. **c.** Representative anisotropy images of stress-induced TDP-35 granules formed in the absence (left) and presence of C-terminally HA-tagged tau (right). Scale bar = 5 µm and 6 µm for without and with tau, respectively. Recruitment of tau-HA was independently confirmed by immunostaining using an anti-HA antibody (see Supplementary Fig. 7d and Methods). **d.** Steady-state anisotropy values extracted from the images in (**c**), for granules formed in the absence and presence of tau. **e.** The apparent homoFRET efficiencies calculated from the corresponding steady-state anisotropy values showing reduced proximity between TDP-35 in the presence of tau. (n = 90 granules measured over 3 independent samples). The data presented indicates the 25^th^ to 75^th^ percentile (box) and 10^th^ to 90^th^ percentile (whiskers). The median (line), mean (white square), and the outliers (black dots) are also shown. All data were statistically significant, as determined by the Mann-Whitney U test (U = 55, p < 0.0001), indicating a significant difference compared with the control.

Arsenite-induced oxidative stress resulted in the formation of cytoplasmic granules by AcGFP-TDP-35 (Supplementary Fig. 7c), which, upon co-expression, colocalized with tau-mCherry as observed by our two-color confocal fluorescence imaging (Fig. 6b). Our *in vitro* cell-free experiments also established that tau and TDP-35 co-phase separate, especially in the presence of RNA (Supplementary Fig. 8), validating our observed colocalization within stress-induced cytoplasmic granules in which RNA is also recruited. We next aimed to further verify the recruitment of TDP-35 and tau in the granules and utilized homoFRET as a nanometric proximity reporter to monitor energy migration between AcGFP-TDP-35. HomoFRET is associated with excited-state energy migration between spectrally identical fluorophores, in which increased molecular proximity enhances energy transfer efficiency, which can be quantified by measuring the loss in the fluorescence anisotropy that arises due to fluorescence depolarization during intermolecular energy hopping within the Förster distance^64,76^. To avoid the complexity introduced by heteroFRET between AcGFP and mCherry, we used fluorescently tagged TDP-35 (AcGFP-TDP-35) and C-terminal HA-tagged tau (devoid of a fluorescent tag) for the homoFRET experiments. We independently confirmed the recruitment of tau-HA into these stress-induced TDP-35 granules (Supplementary Fig. 7d). In the presence of tau, the fluorescence anisotropy of AcGFP-TDP-35 showed a significant increase relative to the TDP-35-only granules, indicating a reduction in the homoFRET efficiency within condensates (Fig. 6c-e). Such a drop in homoFRET indicated a reduced proximity between AcGFP-TDP-35 within granules, due to an increased intermolecular distance between TDP-35 molecules as a result of the recruitment of tau. Collectively, these findings suggested that both TDP-35 and tau are recruited into cytoplasmic granules during oxidative stress.

### Proline-rich and repeat regions of tau promote co-partitioning with TDP-35 into G3BP1-positive stress granules

To dissect the role of different domains of tau in driving granule formation with TDP-35, we expressed the tau variants, Nh2htau (tau_26-230_) and tau_151-391_, with a C-terminal mCherry tag. Both the tau variants were individually expressed alongside AcGFP-TDP-35, and their recruitment into the stress-induced cytoplasmic granules was confirmed by colocalization (Fig. 7a,b). The colocalization analysis revealed a preferential recruitment of tau_151-391_ followed by full-length tau, and Nh2htau (tau_26-230_) (Fig. 7c). This result is in accordance with our *in vitro* cell-free observations (Fig. 3), which highlighted the role of the proline-rich (residues 151-243) and MTBR (residues 244-372) domains of tau in driving heterotypic condensation. To verify the direct interaction-mediated partitioning of tau within cytoplasmic granules, we used AcGFP-TDP-35 and tau-mCherry constructs and performed FLIM-FRET (fluorescence lifetime imaging microscopy-based FRET), which is a sensitive reporter of nanometric proximity between TDP-35 and different tau variants. Energy transfer between AcGFP (donor) and mCherry (acceptor) is associated with a decrease in the donor fluorescence lifetime, which we quantified using FLIM in the presence of different tau variants to derive relative FRET efficiencies within the cytoplasmic granules (Fig. 7d,e). In line with our colocalization data, FRET efficiency followed a similar trend, where tau_151-391_ (∼ 0.34) showed higher FRET efficiency than full-length tau (∼ 0.18), and Nh2htau (tau_26-230_) (∼ 0.13) (Fig. 7e). Since FRET operates at a nanoscopic distance separation (1-10 nm), the occurrence of FRET necessitates close nanometric proximity (within Förster distance of ∼ 5 nm) originating from intermolecular interactions. Therefore, FRET efficiency provides a quantitative readout for the interaction between TDP-35 and tau, and its variants. Our in-cell FLIM-FRET measurements, in accordance with our cell-free *in vitro* results, suggested that tau_151-391_ preferentially recruits within TDP-35 granules, indicating the crucial role of residues 151-391 of tau encompassing proline-rich and MTBR domains in driving heterotypic co-assembly.

**Figure 7:**
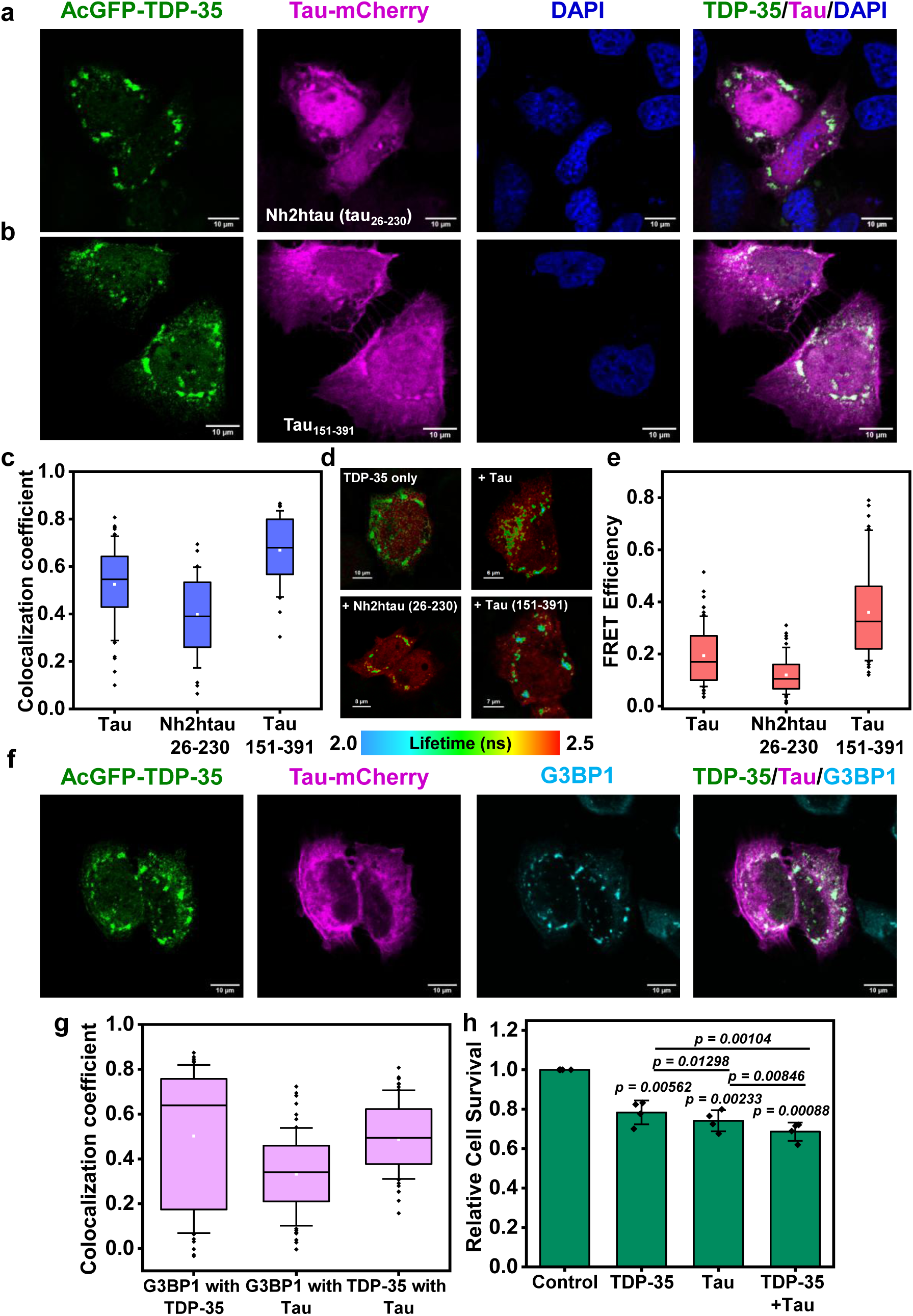
TDP-35:tau granules stain positive for G3BP1. **a.** Representative two-color confocal images of cells overexpressing Nh2htau-mCherry (**a**) and tau_151-391_-mCherry (**b**) with AcGFP-TDP-35. Scale bar = 10 µm. **c.** Two-color colocalization plot quantifying the percent colocalization of AcGFP-TDP-35 with the mCherry-tagged tau variants inside the stress-induced cytoplasmic granules. Data represent mean ± SD (n = 60 (tau) and 45 (Nh2htau and tau trunc) cells measured over 3 independent samples). The p-value was determined using a One-Way ANOVA (p < 0.001), indicating statistical significance among all groups. Analysis for TDP-35:tau was performed on images shown in Fig. 6b and is shown here for comparison. **d.** Representative lifetime images of TDP-35 granules in the absence and presence of different tau variants. **e.** FRET efficiency plot generated from changes in the lifetime of AcGFP-TDP-35 as captured in the lifetime imaging (**d**). The data presented indicates the 25^th^ to 75^th^ percentile (box) and 10^th^ to 90^th^ percentile (whiskers). The median (line), mean (white square), and the outliers (black dots) are also shown. (n = 70 granules measured over 3 independent samples). All data were statistically significant (p < 0.0001) as determined by the Kruskal-Wallis ANOVA test. **f.** Representative confocal images showing colocalization of AcGFP-TDP-35, tau-mCherry, and endogenous G3BP1 (immunostaining) within oxidative stress-induced granules. Scale bar = 10 µm. **g.** Colocalization plot constructed from images shown in (**f**). The data presented indicates the 25^th^ to 75^th^ percentile (box) and 10^th^ to 90^th^ percentile (whiskers). The median (line), mean (white square), and the outliers (black dots) are also shown. (n = 71 cells measured over 3 independent samples) (Data for TDP-35:tau is shown for comparison and is the same as shown in Fig. 5c). **h.** Relative cell survival plot showing co-expression-mediated toxicity under oxidative stress conditions. Data represent mean ± SD (n = 4 independent samples). Statistical significance was determined relative to the control using a paired t-test, and the p-values are reported in the plot.

We next sought to define the nature of these cytoplasmic granules containing both TDP-35 and tau. Given that TDP-43 is known to be associated with stress granules, we asked whether these TDP-35 and tau-positive granules also exhibit features of characteristic stress granules. To this end, we performed immunostaining for G3BP1, a well-established stress granule marker^77^. Oxidative stress-induced TDP-35:tau granules stained positive for G3BP1, confirming their association occurs within *bona fide* stress granules (Fig. 7f, g). Next, to assess the fate and the potential pathological consequences of this association, we aimed to determine the effect of prolonged oxidative stress, which is known to promote aberrant phase transitions into toxic aggregated species. To this end, cells co-expressing TDP-35 and tau were subjected to extended oxidative stress (150 µM sodium arsenite for 4 h), and cell viability was evaluated using the MTT assay. Interestingly, cells co-expressing TDP-35 and tau exhibited greater cytotoxicity, as indicated by the reduced cell survival compared with cells expressing either protein alone (Fig. 7h). This observation suggested that the co-assembly of these two disease-associated proteins within stress granules can exacerbate cellular toxicity under conditions of sustained cellular stress, which is a critical pathological driver of stress granule-driven neurodegeneration. Taken together, our findings demonstrate that stress granules can serve as crucibles for co-phase separation-mediated abnormal phase transitions of TDP-35 and tau, offering a plausible mechanism underlying the exacerbated co-pathology of TDP-43 and tau observed in a range of fatal neurodegenerative diseases.

## Discussion

In this work, we showed heterotypic phase separation of TDP-43 and tau and elucidated the intriguing interplay of molecular drivers of co-condensate formation by dissecting the roles of various domains of both proteins. The summary of our findings is depicted in Fig. 8. Maturation of TDP-43:tau co-condensates via liquid-to-solid phase transitions results in the formation of amyloid-like aggregated species that exhibit cytotoxicity. We also demonstrated that tau and the cytoplasmic variant of TDP-43 co-partition into stress granules formed under oxidative stress conditions. Under extended stress, stress granule-mediated phase transitions of these two proteins can elicit cellular toxicity. *In vitro*, TDP-43 and tau undergo heterotypic phase separation into mesoscopic liquid-like droplets under a carefully chosen condition in which the individual proteins do not undergo homotypic phase separation, indicating the role of intermolecular interactions in co-condensate formation. Our studies revealed the crucial role of intermolecular electrostatic interactions between TDP-43 and tau, whereas hydrophobic interactions play an auxiliary role in driving co-condensate formation. Intriguingly, these TDP-43:tau co-condensates possess dynamic heterogeneity. Our FRAP and FCS measurements indicated that tau is highly mobile within the condensed phase, whereas TDP-43 experiences much slower diffusion in droplets. A critical balance of homo-and heterotypic interactions can potentially dictate the relative mobilities, which, in turn, can define the emergent material properties of TDP-43:tau co-condensates. Slight perturbations in this tightly regulated balance can potentially facilitate molecular rearrangements and rewiring of the interaction network. For instance, RNA alters this interaction network, resulting in enhanced mobility and surface wetting followed by dissolution, indicating an RNA-mediated buffering of the protein phase behavior. This observation recapitulated the cellular role of RNA in stabilizing and solubilizing TDP-43 within the RNA-enriched nucleus^60–62,78^. RNA-dependent phase separation assays revealed competitive interactions between TDP-43, tau, and RNA, supporting the vital role of electrostatics in driving heterotypic phase separation.

**Figure 8:**
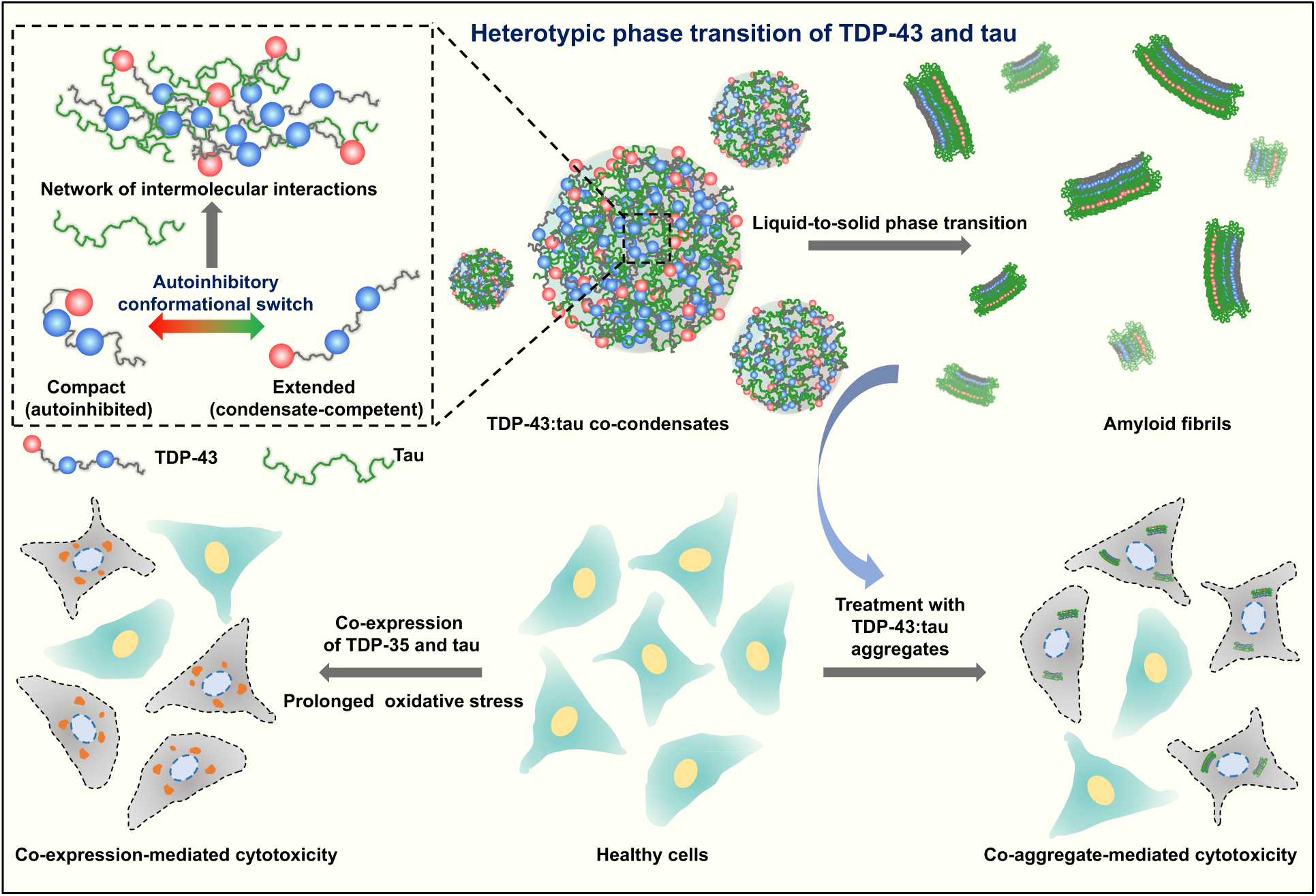
Schematic illustration of heterotypic phase transition of TDP-43 and tau summarizing our findings.

Our findings provide mechanistic underpinnings of heterotypic association-mediated co-condensation and aberrant liquid-to-solid phase transitions. Our results discerned the role of different domains in co-phase separation. The NRR of TDP-43, comprising the NTD and RRMs, and proline-rich and repeat regions of tau are sufficient to drive co-phase separation of TDP-43 and tau. Our single-molecule FRET results unmasked key molecular events associated with co-condensation. We posit that a key conformational switch in TDP-43, from a compact autoinhibited state to an expanded condensate-promoting state, dictates the course of phase separation. Such a structural unwinding event breaks the intramolecular contacts between the NTD and the RRMs and allows TDP-43 to form new intermolecular crosslinks with tau, thereby ensuing co-condensation. The region encompassing the NTD and RRM1 experienced a significant conformational expansion, presumably due to tau-induced weakening of electrostatic intramolecular interactions between oppositely charged NTD and RRM1, and therefore, tipping the balance in favor of intermolecular contacts that promote phase separation. Such an autoinhibitory conformational switch was proposed for a key stress granule-forming protein, G3BP1^77^, suggesting such a regulatory mechanism could play a much broader role in condensate biology.

An interplay between transient intra-and intermolecular interactions governs the material properties of liquid-like condensates that can gradually mature into solid-like aggregates. The aging of TDP-43:tau co-condensates resulted in a transition into β-sheet-rich, amyloid-like fibrils, with kinetics faster than the amyloid conversion of tau. TDP-43 has also been found to be associated with the accumulation of toxic nonfibrillar oligomers of tau^48^ and amyloid-β^79^, thereby exacerbating the AD pathology. In our case, accelerated aggregation of TDP-43:tau suggested that TDP-43 could act as a potent driver when in heterotypic contact with tau in a context dependent-manner, possibly by stabilizing amyloid-competent tau conformations or by providing intermolecular β-sheet templates through the low-complexity domain of TDP-43. These co-aggregates elicited relatively higher cellular toxicity, consistent with clinical observations of worsened neurodegeneration in cases of TDP-43 and tau co-pathologies. Our cellular studies also established enrichment of tau with TDP-35, a disease-associated cleavage product of TDP-43, devoid of the NLS and lacking the normal nucleocytoplasmic shuttling ability. This abnormal cleavage results in the loss of the nuclear TDP-43 pool that can promote further sequestration of full-length TDP-43 within cytoplasmic inclusions via protein and RNA-mediated interactions^75^. Previous reports have also indicated independent recruitment of both TDP-43 and tau into stress granules, where tau associates predominantly with TIA-1-positive granules^63^, whereas TDP-43 can be recruited into either G3BP1-or TIA-1-marked granules^30,38,77^. Our results showed that oxidative stress can lead to co-partitioning of TDP-35 and tau in G3BP1-positive stress granules in a domain-specific manner. Such stress granules can potentially serve as crucibles for pathological aggregation^29,30^. Consistent with this hypothesis, we observed that under prolonged stress, cells with TDP-35 and tau-containing stress granules displayed considerable cellular toxicity. Therefore, cellular crosstalk between TDP-43 and tau can offer a crucial molecular basis for disease-associated synergism and co-deposition found in a wide range of neuropathological conditions. The growing evidence of a complex interplay of multiple proteins contributing to more severe forms of pre-existing neuropathological conditions has opened new unexpected avenues^47–51,80^, including highlighting the roles of posttranslational modifications^81^ and newfound short RNA chaperones^78^ in regulating TDP-43 phase separation and aggregation. These emerging discoveries, together with our findings, will provide a broad mechanistic framework for phase transition-mediated disease progression and pave the way for therapeutic interventions by regulating heterotypic interactions and tuning the physical properties of biomolecular condensates.

## Methods

### Construct details

Bacterial expression constructs of TDP-43-MBP in pJ4M (Addgene Plasmid #104480, https://www.addgene.org/104480/) and TDP-43 CTD in pJ411 (Addgene Plasmid #98669, https://www.addgene.org/98669/) were a kind gift from Nicolas Fawzi. The N-terminal truncation (NRR, residues 1-259, cloned in pJ411) and other point mutations were created from the full-length TDP-43-MBP construct. A single-cysteine variant of TDP-43 (with 244C intact) was synthesized by GeneToProtein/Biotech Desk. and used to create the null-cysteine, single-cysteine, and dual-cysteine FRET constructs (wild-type and monomerization variants) via site-directed mutagenesis (using KOD Hot Start Master Mix, Cat. No. 71842, Sigma-Aldrich). Mammalian expression constructs for TDP-43 and TDP-35 were cloned from HeLa cDNA into the pAcGFPC1 vector. All tau constructs used in this study were created from a 6×His-Tau 2N4R-17C plasmid, gifted to us by Elizabeth Rhoades (University of Pennsylvania, USA), as described previously^26^. After removal of the 6×His tag via cloning, all cysteine variants and PHF6* (Δ275-280) and PHF6* (Δ306-311) deletion mutants were generated by site-directed mutagenesis using a QuickChange kit (Stratagene). The phosphomimetic variant used in this study (tau 4E MARK) was synthesized by GeneToProtein/Biotech Desk. The mammalian full-length construct was created by cloning tau from the tag-less construct separately into pmCherryN1 and pcDNA 3.1(-) for a C-terminal HA tag. The bacterial and mammalian expression domain-specific variants, including tau [Nh2htau and tau_151-391_], were created via cloning into pET11a or pmCherryN1, respectively. Unless mentioned, all *in vitro* experiments in this study were performed using a null-cysteine variant of tau (C291S, C322S). All the mutations were verified using sequencing. See Table S1 for more details on the constructs and primers used in this study.

### Recombinant protein expression and purification

For recombinant expression of TDP-43, all plasmids were transformed into *E. coli.* BL21 (DE3) standard cells grown in LB media (37 °C, 220 rpm). For full-length TDP-43 and mutants, cells were grown to an O.D._600_ of 0.6-0.8, and overexpression was induced with 1 mM isopropyl-β-thiogalactopyranoside (IPTG) for 16 h at 16 °C. Bacterial cultures were pelleted by centrifugation at 3220 × g at 4 °C and stored at-80 °C for further purification. Wild-type and mutant TDP-43 pellets were resuspended in buffer (20 mM Tris-Cl, 1 M NaCl, 10 mM imidazole, 1 mM phenylmethylsulfonyl fluoride (PMSF), 4 mM β-mercaptoethanol (BME) with 10 % glycerol, pH 8.0) for cell lysis using probe sonication (5 % amplitude, 15 s ON, and 10 s OFF) for 25 min. The cell debris was separated by centrifugation at 15,557 × g, 4 °C for 1 h, followed by binding of the supernatant onto a pre-equilibrated Ni-NTA column. The bound protein was washed with buffer (20 mM Tris-Cl, 1 M NaCl, 10 mM imidazole, and 4 mM BME, pH 8.0) and eluted using 300 mM imidazole. The Ni-NTA eluate was then concentrated using 50 kDa MWCO Amicon filters before loading onto a HiLoad 16/600 Superdex G-200 (GE) for size-exclusion chromatography (SEC). Additional impurities were further removed by SEC, and the protein was buffer-exchanged into storage buffer (20 mM Tris-Cl, 300 mM NaCl, 1 mM dithiothreitol (DTT)). The eluted fractions were run on SDS-PAGE to assess purity, after which the pure protein fractions were pooled and concentrated using 50 kDa MWCO Amicon filters. The protein concentration was estimated by measuring absorbance at 280 nm (ε_280_ = 1,12,020 M^−1^ cm^−1^) and stored at-80 °C for future use.

For TDP-43 NRR-pJ411 (residues 1-259), cells were grown to an O.D._600_ of ∼0.8, and protein expression was induced with 0.5 mM IPTG for 16 h at 16 °C. The cell pellets were thawed at room temperature and resuspended in lysis buffer (20 mM Tris base, 300 mM NaCl, 10 mM imidazole, 4 mM BME, with 10 % glycerol, pH 8). Cell lysis was performed using a probe sonicator (5 % amplitude, 15 s ON, 10 s OFF) for 20 min, and the supernatant was separated by centrifugation at 15,557 × g, 4 °C for 1 h. The soluble protein was loaded onto a Ni-NTA column, the non-specifically bound proteins were removed by washing, and the protein of interest was eluted in the Ni-NTA elution buffer (20 mM Tris base, 300 mM NaCl, 300 mM imidazole, 1 mM DTT). For removal of the C-terminal 6×His tag, the eluted protein was incubated with Tobacco Etch Virus (TEV) protease at a molar ratio of 1:30 (TEV:protein) and kept at room temperature for 1.5 h following dialysis into an imidazole-free buffer using a 3 kDa MWCO membrane at 4 °C overnight. The dialyzed protein was further passed through a recharged Ni-NTA column to separate the pure, cleaved protein, which was then concentrated using 10 kDa MWCO Amicon filters and stored at-80 °C for future use. Protein purity was assessed by running the samples on an SDS-PAGE gel, and the concentration was estimated by measuring absorbance at 280 nm (ε_280_ = 26,030 M^−1^ cm^−1^).

The C-terminal domain of TDP-43 (267-414) expression was induced at O.D._600_ of ∼ 0.8 with 1 mM IPTG at 37 °C for 4 h. The protein was purified under denaturing conditions and later buffer-exchanged into native conditions prior to phase separation studies. The cell pellet was resuspended in lysis buffer (20 mM Tris base, 500 mM NaCl, 10 mM imidazole, 4 mM BME, pH 8.0) and lysed to obtain inclusion bodies using probe sonication (10 % amplitude, 15 s ON, 10 s OFF) for 25 min. The supernatant was discarded after centrifugation, and the inclusion bodies were further dissolved in the denaturation buffer (8 M Urea, 20 mM Tris base, 500 mM NaCl, 10 mM imidazole, 4 mM BME, pH 8.0) with intermittent mixing. The dissolved lysate was then centrifuged at 15,557 × g, and the supernatant was loaded on pre-equilibrated Ni-NTA beads. The column was then washed with denaturation buffer, and the protein was eluted with 300 mM imidazole. Next, the protein was buffer-exchanged into a native buffer (20 mM Tris base, 500 mM guanidium hydrochloride, pH 8.0) to facilitate TEV-mediated cleavage of the C-terminal 6×His tag. The protein was incubated with TEV protease (molar ratio 1:20) and kept overnight at room temperature. The cleaved protein was passed through a freshly recharged column, and the flow-through was collected and denatured by adding 8 M urea directly to the protein. The denatured protein was concentrated using 3 kDa MWCO Amicon filters and buffer-exchanged into the storage buffer (8 M Urea, 20 mM Tris base, pH 8.0) using a PD-10 column, and stored at-80 °C until further use. Pure protein was buffer-exchanged using a NAP-5 column into 10 mM MES buffer, pH 5.5, for phase separation studies. Protein purity was assessed by running the samples on an SDS-PAGE gel, and the concentration was estimated by measuring absorbance at 280 nm (ε_280_ = 18,350 M^−1^ cm^−1^).

For wild-type and other variants of tau, plasmids were expressed in *E. coli.* BL21 (DE3) standard cells were grown till O.D._600_ of 0.6 in LB media (220 rpm, 37 °C), and induction was done using 0.5 mM IPTG at 37 °C for 1 h. The proteins were purified as described previously^31^. In brief, the cell pellets were resuspended in lysis buffer (20 mM MES, 500 mM NaCl, 1 mM MgCl2, 1 mM EDTA, 1 mM PMSF, and 2 mM DTT, pH 6.5) and subjected to probe sonication (5 % amplitude, 15 s ON, 10 s OFF) for 25 min. The supernatant was separated by centrifugation and incubated with streptomycin sulfate and glacial acetic acid to remove bound nucleic acids, followed by salting out the protein with 60 % ammonium sulfate. The precipitated protein was extracted by centrifugation at 15,557 × g at 4 °C. The pellets were further dissolved and loaded onto an SP-Sepharose column (for tau and tau_151-391_ variant) and a Q-Sepharose (for Nh2htau) in a low-salt buffer. The proteins were then eluted by a linear salt gradient of high-salt buffer (20 mM MES, 1 M NaCl, 1 mM EDTA, 2 mM DTT, 1 mM MgCl2, and 1 mM PMSF, pH 6.5). The eluted fractions were run on a gel, pooled, and further purified and buffer exchanged into storage buffer (25 mM HEPES, 50 mM NaCl, 2 mM DTT, pH 7.4) using a HiLoad 16/600 Superdex G-200 (GE) column for size-exclusion chromatography (SEC). The SEC fractions containing protein of interest were pooled, concentrated using 10 kDa MWCO Amicon filters, and the concentration was estimated by measuring absorbance at 280 nm before storage in-80 °C for future use. [ε_280_ = 6400 M^−1^ cm^−1^ (tau and tau_Δ275-280_), ε_280_ = 5120 M^−1^ cm^−1^ (tau_Δ306-311_), and ε_280_ = 2560 M^−1^ cm^−1^ (tau_151-391_ and Nh2htau)]. The single-cysteine variants of full-length tau were buffer exchanged into denaturing conditions (6 M GdMCl, 10 mM HEPES, 50 mM NaCl, 0.3 mM tris(2-carboxyethyl)phosphine (TCEP), pH 7.4) during SEC for further labeling reactions.

### Fluorescence labeling

TDP-43 single-cysteine variants were site-specifically labeled with fluorescein-5-maleimide (F5M) using the thiol-maleimide chemistry for fluorescence anisotropy studies. Freshly thawed proteins were incubated with 0.3 mM TCEP under native conditions (20 mM Tris-Cl, 300 mM NaCl, pH 7.4) and kept on ice for 45 min to ensure complete reduction of cysteine residues prior to labeling. F5M dye was later added to the protein in a molar ratio of 1:20 (protein:dye), and the reaction was incubated at 25 °C, under shaking conditions for 3.5 h. TDP-43 244C (single-cysteine) and truncation variants were labeled under similar conditions with AlexaFluor488-maleimide (molar ratio 1:3; protein:dye) for imaging, FRAP, and FCS measurements. For single-molecule FRET measurements, the dual-labeled cysteine variants of TDP-43 were stochastically labeled with AlexaFluor488-and AlexaFluor594-maleimide as donor and acceptor fluorophores, respectively. TCEP-treated proteins were incubated with the donor dye at a 1:0.9 (protein:dye) molar ratio under shaking conditions for 1.5 h. The acceptor dye was later added to the same reaction mixture at a molar ratio of 1:4.5 (protein:dye), and the mixture was incubated at room temperature in the dark for 5 h. Excess, unbound dye was removed by washing, and the labeled protein was concentrated using 50 kDa MWCO Amicon filters at 4 °C.

For tau, single-cysteine and truncation variants were labeled with AlexaFluor594-maleimide, AlexaFluor488-maleimide, and F5M for imaging, FCS, and steady-state anisotropy experiments, respectively. The pure protein stored under denaturing conditions was mixed with Alexa dyes (molar ratio 1:3) or F5M dye (molar ratio 1:20), and kept in the dark under stirring conditions for 4 h at room temperature. The removal of free dye was performed using 10 kDa MWCO Amicon filters, and the labeled protein was gradually buffer-exchanged into native storage buffer (25 mM HEPES, 50 mM NaCl, 2 mM DTT, pH 7.4). The labeling efficiency and labeled protein concentrations were estimated by measuring the absorbance at 280 nm for proteins (see above), 494 nm for AlexaFluor488 (ε_494_ = 73,000 M^−1^ cm^−1^), and F5M (ε_494_ = 68,000 M^−1^ cm^−1^), and 590 nm for AlexaFluor594 (ε_494_ = 92,000 M^−1^ cm^−1^).

### Phase separation and turbidity assays

Phase separation of TDP-43 and tau was induced by cleaving the TDP-43 C-terminal solubilizing MBP-6×His tag with TEV protease at a molar ratio of 1:10 (TEV:protein) in 20 mM HEPES, 15 mM NaCl, 1 mM DTT, pH 7.4. TDP-43 concentration was fixed at 5 µM (unless mentioned otherwise), and tau was varied from 0 to 30 µM for the stoichiometry-based assay. Truncated TDP-43 (NRR, 5 µM, and CTD, 10 µM) were mixed with tau (20 µM) and incubated at room temperature for 10 min. For tau variants (Nh2htau and tau_151-391_, 20 µM), TDP-43 (5 µM) was incubated with TEV protease (1:10, TEV:protein) in the presence of tau at room temperature for 10 min. To probe the role of hydrophobic interactions and the effect of RNA on heterotypic TDP-43:tau condensates, proteins were mixed at a fixed molar ratio of 1:4 (5 µM TDP-43 and 20 µM tau) in the presence of increasing concentrations of hexanediol and polyU RNA. For salt-dependent phase separation under physiological conditions, reactions were set up in the presence of 10 % PEG 8000 with added NaCl concentrations (0-300 mM). For turbidity measurements, the absorbance of TDP-43:tau phase separation reactions at 350 nm was monitored under various conditions. All reactions were incubated at room temperature for 10 min prior to measurement. For the stoichiometry-dependent turbidity assays and to study the effect of different mutants of TDP-43, the proteins were mixed in the required ratios while keeping the salt concentration constant (15 mM NaCl). For all reactions, absorbance was measured using a Thermo Fisher Scientific NanoDrop OneC Spectrophotometer. All the data were repeated independently three times and are plotted as mean ± SD.

### Confocal microscopy and image analysis

Confocal fluorescence imaging of droplets was performed on a Zeiss LSM 980 Elyra 7 super-resolution microscope, recorded with an oil-immersion 63× objective (NA 1.4), and detected with a high-resolution monochrome cooled AxioCamMRm Rev. 3 FireWire(D) camera. Phase separation reactions with the full-length or the truncated variants (as mentioned) were set up in 20 mM HEPES, 1 mM DTT, pH 7.4 buffer, and doped with 1 % of the corresponding AlexaFluor488-(TDP-43) and AlexaFluor594-(tau) labeled proteins for visualization. The droplet formation was induced with TEV protease (1:10; TEV:protein). All the reactions were incubated at room temperature for 10 min before dropcasting 5-10 µL of the reaction on a glass coverslip. Droplets were allowed to settle for 2-3 min and imaged with a 590 nm excitation and a 488 nm laser diode (11.9 mW) for two-color imaging of the heterotypic condensates. The emitted fluorescence was recorded by the Airyscan 2 detector equipped with 32 channels (GaAsP) in the confocal laser scanning mode to obtain images with 1840×1840-pixel per region and 16-bit depth. Airyscan processing was performed using the built-in Zen Blue 3.2 software. The colocalization and partition coefficient analyses were performed in ImageJ (NIH, Bethesda, USA). Pearson’s and Manders’ colocalization coefficient was estimated for the 1:4 stoichiometry (5 µM TDP-43 and 20 µM tau) using the JACoP plugin in ImageJ software. For partition coefficient (K_app_) determination, background-corrected intensities for each channel (green and red) were measured separately in the condensate (I_dense_) and dispersed phase (I_light_) to calculate the apparent partition coefficient (using Equation 1) as a function of varying tau concentrations in the phase separation reactions.

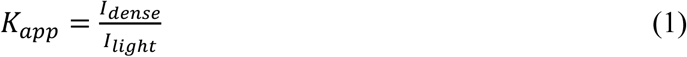

### Fluorescence recovery after photobleaching (FRAP) measurements

The FRAP experiments were done on a ZEISS LSM980 confocal microscope. All condensate samples, doped with 1 % AlexaFluor488-labeled protein, were focused using a 63× oil-immersion objective lens (NA 1.4). A 0.7 µm-diameter area was photobleached using a 488 nm laser at 100 % power, and subsequent recovery of fluorescence intensity was monitored using Zen Blue 3.2 software. For every condition, a minimum of 7 recovery profiles from independent condensates were acquired. The normalized recovery profiles were background-subtracted and plotted in Origin software.

### Fluorescence correlation spectroscopy (FCS) measurements

All FCS measurements were performed on a MicroTime 200, PicoQuant time-resolved confocal microscope. Briefly, the system was calibrated using a 10 nM solution of AlexaFluor488 dye to determine the confocal volume (V_eff_, 0.54 fL) and the structure parameter (κ, 4.82). These parameters were used to fit data recorded for the dispersed phase and within droplets. FCS measurements were performed by separately doping AlexaFluor488-labeled single-cysteine variants of TDP-43 and tau in the dispersed (15-20 nM) and droplet (5-10 nM) phase. Freshly prepared samples (30 µL) were drop-cast, and data were recorded 50 µm within the solution for the dispersed phase, or via focusing individual droplets for phase-separated samples. The recorded data were fit using a triplet state model as follows:

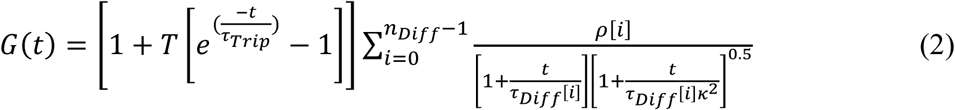

Here, G(t), ρ, T, τ_Trip_, τ_Diff_, and κ denote the correlation amplitude, contribution of the i^th^ diffusing species, fraction of the triplet state, lifetime of the triplet state, diffusion time of the i^th^ diffusing species, and the structure parameter of the corresponding confocal volume, respectively.

### Circular dichroism (CD) spectroscopy

Far-UV CD spectra were acquired for the purified TDP-43 and its null-cysteine variant proteins. The measurements were performed on a Chirascan spectrophotometer (Applied Photophysics, UK) using a 1 mm path-length quartz cuvette. Each purified protein was diluted to 3 µM in 5 mM sodium phosphate buffer with 5 mM NaCl. A total of 10 scans were taken for each sample, which were further averaged and blank-subtracted using the ProData Viewer software (ver. 4.1.9) and plotted using the Origin software.

### Single-molecule FRET measurements

Single-molecule FRET measurements were performed by the ratiometric detection of donor and acceptor emission intensities in pulsed-interleaved excitation (PIE) mode using a confocal-based time-resolved microscope (MicroTime 200, PicoQuant) as described previously^26,68^. All the measurements were performed in the 20 mM HEPES, 1 mM DTT, pH 7.4 buffer supplemented with n-propyl gallate as an oxygen scavenger. For the dispersed phase measurements, dual-labeled TDP-43 was incubated with TEV protease to cleave the C-terminal MBP-6×His tag and measurements were performed in the presence of 100-200 pM of labeled protein. For the droplet phase measurements, TDP-43 (5 µM) and tau (20 µM) were phase separated in the presence of 25-30 pM of dual-labeled TDP-43, and the reactions were incubated at room temperature for 10 min before dropcasting onto the glass coverslip. The samples were excited sequentially with the donor (485 nm) and acceptor (594 nm) pulsed lasers with a repetition rate of 20 MHz using a two-color pulsed-interleaved excitation (PIE) mode, with the total laser power fixed at 50-60 µW (for dispersed phase) and 6-8 µW (for droplet phase), as measured at the back aperture of the objective. The emitted photos were collected from ∼ 50 µm (for dispersed phase) or ∼ 2-3 µm (for condensates) inside the coverslip surface by a 60× water-immersion objective (NA 1.2, Olympus). The fluorescence was further filtered with a dichroic 485/594 nm filter and a 500 nm longpass filter, then passed through a 50 µm pinhole to remove out-of-plane emission. The emitted photons were separated by a dichroic beamsplitter (zt594rdc) and further filtered with 530/50 nm and 645/75 nm bandpass filters for the donor and acceptor channels, respectively. The filtered photons were then separately detected by two single-photon avalanche diodes (SPADs) and used for ratiometric PIE-FRET calculations with the in-built SymphoTime64 software v2.7.

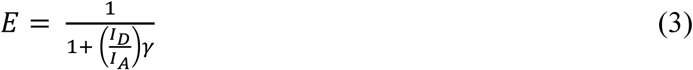

where 𝐼_𝐷_and 𝐼_𝐴_are the donor and acceptor intensities, which are corrected for the different detector efficiencies (donor and acceptor channel) and fluorophore quantum yields (𝛾). The system calculated the background, and a binning time of 0.5 ms and a PIE threshold of 35 counts were applied to select bursts containing donor and acceptor fluorophores. The PIE-FRET histogram was constructed after correcting for the differential detector efficiencies among the channels (γ = 1.08 and 0.71 for the 39C-144C and 144C-244C constructs), the fraction of acceptor molecules directly excited by the donor laser (β = 0.003), and lastly to consider the donor-acceptor spectral crosstalk (α = 0.05). The PIE-FRET mode eliminated zero-FRET events that arise due to donor-only and acceptor photobleaching events. The total number of events collected for both constructs was at least 20,000. The plotting and Gaussian fitting of the combined FRET histograms were performed using Origin.

### Aggregation kinetics

Aggregation reactions were monitored by recording the fluorescence kinetics of an amyloid-marker, ThT, on a POLARstar Omega Plate Reader Spectrophotometer (BMG LABTECH, Germany). Aggregation reactions (150 µL) were set up under physiological conditions (20 mM HEPES, 150 mM NaCl, 10 % PEG 8000, 1 mM DTT, pH 7.4) for both homotypic and heterotypic phase-separated TDP-43 (5 µM), tau (20 µM), and TDP-43 (5 µM) + tau (20 µM). ThT was added at a concentration of 20 µM, and the reactions were set up in triplicate in 96-well plates. A glass bead (3-mm diameter) was added to each well to facilitate mixing, and the reactions were incubated at 25 °C under stirring conditions (300 rpm) for 16 h. Time-dependent fluorescence intensity was measured by exciting the samples at 485 nm, and the data were plotted in Origin.

### Transmission electron microscopy

Following saturation of the aggregation reactions (16 h for homotypic aggregates and 10 h for heterotypic aggregates), the samples were centrifuged at 25,000 × g for 1 h, and the pellets were resuspended in 10 µL distilled water. For sample preparation, 3 µL of the resuspension was dropcast onto a 300-mesh carbon-coated electron microscopy grid, allowed to bind for 2 min at room temperature. The excess sample was soaked off, and 3 µL of the staining solution (1 % w/v uranyl acetate) was incubated for 2 min, followed by soaking off the excess solution. The grids were dried overnight at room temperature, and images were obtained using a JEOL JEM-200.

### Vibrational Raman spectroscopy

Vibrational Raman spectra were obtained for the homotypic and heterotypic aggregates formed upon aging of the condensates under physiological conditions. Aggregation reactions were centrifuged at 25,000 × g for 1 h, following saturation in the ThT kinetics at 10 h (for TDP-43:tau heterotypic aggregates) and 16 h (TDP-43 and tau homotypic aggregates). The pellet was washed multiple times with distilled water and resuspended in 10 µL of water. The measurements were performed at room temperature on an inVia laser Raman microscope (Renishaw, UK). The resuspended aggregates (5 µL) were dropcast onto aluminum foil-covered glass slides and focused with a 100× long working distance objective (NA 0.85, Leica). For excitation, a near-infrared laser (785 nm) was used at 100 % power (500 mW) with a 10-second exposure time, following which the scattered photons were filtered out by an edge filter blocking the Rayleigh scattering. The Raman scattering was further dispersed by passing through a diffraction grating (1200 lines/mm) and detected by an air-cooled CCD detector. All spectra were collected over 10 accumulations and subjected to baseline correction and smoothening using the Wire 3.4 software. The data were normalized with respect to the phenylalanine peak (∼ 1003 cm^-1^) for plotting and analysis using the Origin software. The secondary structural component analysis was performed by Gaussian fitting and deconvolution of the Amide peaks in Origin.

### Mammalian cell culture

HeLa cells were maintained in Dulbecco’s Modified Eagle’s Medium (DMEM) supplemented with 10 % heat-inactivated fetal bovine serum (FBS), 100 U/mL penicillin, and 100 µg/mL streptomycin at 37 °C in a humidified incubator with 5 % CO₂. For transient overexpression, cells were seeded on glass coverslips, transfected with 500 ng of plasmid DNA using X-tremeGENE HP DNA transfection reagent (Roche), and grown for 22 h. To recapitulate oxidative stress conditions, cells were treated with 300 µM sodium arsenite for 1 h, then fixed with 4 % paraformaldehyde (PFA) in PHEM buffer (60 mM PIPES, 25 mM HEPES, 10 mM EGTA, 2 mM MgCl2, pH 6.8) for 10 min. For visualization, the nuclei were stained with DAPI before mounting on slides with Fluoromount-G (Southern Biotech).

### Immunostaining

HeLa cells transfected with the plasmids of interest and grown on glass coverslips for 22 h were fixed with 4 % PFA in PHEM buffer. Cells were permeabilized with 0.2 % saponin and incubated with primary antibodies [Anti-HA tag antibody (clone 16B12, mouse monoclonal (dilution 1:500), BioLegend, Cat. No. 901503) and Anti-G3BP1 antibody (rabbit polyclonal (dilution 1:500), Proteintech, Rosemont, IL, USA; Cat. No. 13057-2-AP)] diluted in blocking buffer (5 % FBS in PHEM) and kept overnight at 4°C in a humidified chamber. Subsequently, the coverslips were washed with PBS and incubated with fluorophore-conjugated secondary antibodies (AlexaFluor568 Goat anti-mouse IgG, Cat. No. A11031 for tau-HA and AlexaFluor405 Goat anti-rabbit IgG, Cat. No. A31556 for G3BP1; 1:500 dilution) in the dark at room temperature for 1 h. The coverslips were then washed with PBS and mounted on glass slides using Fluoromount-G.

### Confocal imaging and colocalization analysis

Confocal imaging of fixed cells was performed using a Zeiss 710 confocal laser scanning microscope with a 63× oil-immersion lens (NA 1.4). Samples were excited at 405 nm (DAPI/G3BP1), 488 nm (AcGFP), and 561 nm (mCherry/tau-HA), and imaging was performed with a 1024 × 1024-pixel-per-image and 8-bit depth using ZEN 2012 v. 8.0.1.273 software (ZEISS). Image processing of representative confocal images and co-localization analysis were performed using ImageJ (NIH, Bethesda, USA). For colocalization analysis, cytoplasmic granules were selected using automatic thresholding, and Pearson’s correlation coefficient (r) values were obtained for individual cells using the JACoP plugin. Data were collected across three independent experiments and plotted to represent the distribution of colocalization for various protein pairs.

### MTT-based cell viability assay

Cell viability was assessed using the MTT (3-(4,5-dimethylthiazol-2-yl)-2,5-diphenyltetrazolium bromide) colorimetric assay (EZcountTM MTT Cell Assay Kit, HiMedia, Cat. No. CCK003-1000). For aggregate-based MTT, HeLa cells were incubated with 5 µg of aggregates (TDP-43 alone, tau alone, or TDP-43:tau aggregates) and seeded at a density of 10^4^ cells per well in a 96-well plate. The cells were grown at 37 °C with 5 % CO_2_ for 72 h post-treatment, and then 10 µL MTT reagent was added to each well. The formazan crystals were allowed to form for 4 h and then dissolved in a solubilization buffer by mixing. The formation of crystals was quantified by measuring the absorbance at 570 nm with a reference wavelength of 630 nm on a POLARstar Omega Plate Reader Spectrophotometer (BMG LABTECH, Germany). Similarly, for the expression-based MTT, 4 sets of cells were transfected: pAcGFPC1 with pmCherryN1 (vector control), pAcGFP-TDP-35 with pmCherryN1 (TDP-35 control), pAcGFPC1 with tau-pmCherryN1(tau control), and pAcGFP-TDP-35 with tau-pmCherryN1 (TDP-35 + tau). Post-transfection (24 h), 10^4^ cells were seeded per well in a 96-well plate and allowed to adhere for 4 h. The MTT reagent (10 µL) was then added to each well, and the mixture was incubated at 37°C for 4 h. Cell viability was quantified by measuring the absorbance of dissolved crystals at 570 nm and plotted as a normalized value relative to the empty vector control. All treatments were performed in triplicate and repeated across at least three independent experiments.

### Fluorescence anisotropy and homoFRET imaging

Steady-state fluorescence anisotropy imaging was performed on the MicroTime 200 time-resolved confocal microscope (PicoQuant, Germany). For the dispersed phase measurements, single-cysteine F5M-labeled variants of TDP-43 (39, 144, 244, 333, and 403) and tau (199, 244, 322, and 400) were diluted to 150-200 nM concentration. For measurements inside droplets, TDP-43 (5 µM) and tau (20 µM) were mixed, and phase separation was induced with TEV (1:10 molar ratio) in the presence of 15-30 nM F5M-labeled variants. The samples were excited with a 485 nm laser using a 60× Super Apochromat water-immersion objective (NA 1.2, Olympus). The emission was passed through a 485/594 dichroic mirror followed by filtering by a 530/50 nm bandpass. The out-of-focus light is removed using a 50 μm pinhole, and the donor emission is split into two channels (parallel and perpendicular) using a beamsplit polarizer, with detection on separate single-photon avalanche diodes (SPADs). Dispersed phase and single-droplet anisotropy values were calculated after incorporating the lens correction factors using the following equation in the in-built SymphoTime64 software v2.7.

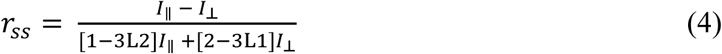

where 𝐼_∥_ and 𝐼_⊥_are the parallel and perpendicular intensities and L1 (0.0308) and L2 (0.0368) are the objective lens correction factors^82^. For homoFRET imaging, cells expressing AcGFP-TDP-35 alone and with tau-HA were fixed after 1 h of sodium arsenite treatment and mounted using Fluoromount-G. The anisotropy values in the absence (𝑟_𝑠𝑠0_) and presence (𝑟_𝑠𝑠_) of homoFRET were used to estimate the apparent uncorrected homoFRET efficiencies using equation (5). Cytoplasmically dispersed TDP-35 was used as the non-homoFRET control for estimating the E_homo_.

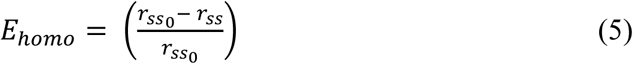

### FLIM-FRET measurements

FLIM-FRET was performed on the above-mentioned time-resolved microscope (MicroTime 200, PicoQuant, Germany) equipped with picosecond lasers and single-photon detection. HeLa cells expressing AcGFP-TDP-35 alone and with different tau-mCherry variants were treated with 300 µM sodium arsenite for 1 h to facilitate stress granule formation, then fixed with 4 % PFA. The samples were excited with a pulsed laser (485 nm) and imaged with a water-immersion Super Apochromat 60× objective (NA 1.2; Olympus). The imaging was performed with a 10 µs dwell time and 256 × 256 pixels per region of interest. The emitted photons were collected, passed through a 530/50 nm bandpass, filtered by a pinhole (50 µm) to obtain in-focus images, and detected by the SPADs in time-correlated single-photon counting (TCSPC) mode. The donor lifetime was calculated by fitting the decay of individual TDP-35 granules formed in the absence of an acceptor (mCherry) using equation (6), in SymphoTime64 software v2.7.

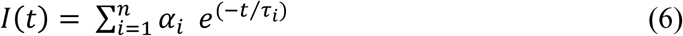

where 𝜏_𝑖_ are the individual decay times with their respective amplitudes (𝛼_𝑖_) integrated over the n-number of exponents. The donor decay was fitted using a bi-exponential reconvolution model with corrections for the calculated background and IRF shift, with parameter optimization performed using a maximum likelihood estimator (MLE). The donor lifetime in the presence of various tau variants tagged with mCherry was recorded, and lifetime-based FRET efficiencies (E) were calculated by equation (7) using the commercially available SymphoTime64 software v2.7,

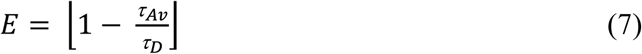

where, *τ*_𝐴𝑣_ is the amplitude-average lifetime of the donor in the presence of the acceptor and *τ*_𝐷_ (2.08 ns) is the unquenched donor lifetime in the absence of FRET.

### Statistics and reproducibility

All experiments were repeated with at least three independent samples; the exact n is provided in the figure legends. Data plotting and analyses were performed using the Origin software. Statistical analyses were performed using a paired t-test for the MTT assay and a One-Way ANOVA or a Mann-Whitney U/Kruskal-Wallis test for larger sample sizes. The p-values are mentioned in the respective figures or corresponding legends. The data are presented as mean ± SD, or the windows are specified separately in the box-and-whisker plot legends. Fluorescence microscopy imaging was repeated at least three times for biological replicates, and the analysis software used is described in the Methods or figure legends.

## Acknowledgments

We thank IISER Mohali and Department of Science and Technology, Govt. of India (FIST grant # SR/FST/LS-II/2017/97 to the Department of Biological Sciences, IISER Mohali). We thank Anusandhan National Research Foundation (ANRF) (SUPRA SPR/2020/000333) (J.C. Bose Fellowship JCB/2023/000016 to S.M.), (CRG/2021/002314) and Indo-French Centre for the Promotion of Advanced Research (IFCPAR/CEFIPRA) (IFC/A/6903-3/2023/680) to S.M. for financial support. We acknowledge the Ministry of Education, Govt. of India (STARS-24-0039 to S.M.), Prime Minister’s Research Fellowships (to A.W. and R.K.), and ANRF National Post-doctoral Fellowship (PDF/2025/000065 to A.S.) for financial support. We are grateful to Prof. Mahak Sharma (IISER Mohali) and her lab members, Biswaheree Mahananda and Anamika Kutrial, for providing us with cell culture materials and facilities. We thank Dr. Sandeep Rai for his valuable suggestions, Sneha Bisht and Ayush Gupta for their help with our experiments, and the members of the Mukhopadhyay lab for critically reading the manuscript.

## Author contributions

A.W. and S.M. conceived the project. A.W., S.S., R.K., A.S., D.D.R., and A.J. performed the experiments and analyses. A.W. prepared the figures and wrote the first draft. S.M. supervised the work, wrote/edited the manuscript, obtained funding, and provided the overall direction. All authors discussed the results and commented on the manuscript.

