## Supplementary Information for "An autoinhibitory regulatory switch governs heterotypic phase separation and liquid-to-solid phase transition of TDP-43 and tau into cytotoxic amyloids"

**a** **TDP-43**

```
MSEYIRVTED ENDEPIEIPS EDDGTVLLST VTAQFPGACG LRYRNPVSQC MRGVRLVEGI  60
LHAPDAGWGN LVYVVNYPKD NKRKMDETDA SSAVKVKRAV QKTSDLIVLG LPWKTTEQDL 120
KEYFSTFGEV LMVQVKKDLK TGHSKGFGFV RFTEYETQVK VMSQRHMIDG RWCDCKLPNS 180
KQSQDEPLRS RKVFVGRCTE DMTEDELREF FSQYGDVMDV FIPKPFRAFA FVTFADDQIA 240
QSLCGEDLII KGISVHISNA EPKHNSNRQL ERSGRFGGNP GGFGNQGGFG NSRGGGAGLG 300
NNQGSNMGGG MNFGAFSINP AMMAAAQAAL QSSWGMMGML ASQQNQSGPS GNNQNQGNMQ 360
REPNQAFGSG NNSYSGSNSG AAIGWGSASN AGSGSGFNGG FGSSMDSKSS GWGM       414
```

**b** **Tau**

```
MAEPRQEFEV MEDHAGTYGL GDRKDQGGYT MHQDQEGDTD AGLKESPLQT PTEDGSEEPG  60
SETSDAKSTP TAEDVTAPLV DEGAPGKQAA AQPHTEIPEG TTAEEAGIGD TPSLEDEAAG 120
HVTQARMVSK SKDGTGSDDK KAKGADGKTK IATPRGAAPP GQKGQANATR IPAKTPPAPK 180
TPPSSGEPPK SGDRSGYSSP GSPGTPGSRS RTPSLPTPPT REPKKVAVVR TPPKSPSSAK 240
SRLQTAPVPM PDLKNVKSKI GSTENLKHQP GGGKVQIINK KLDLSNVQSK CGSKDNIKHV 300
PGGGSVQIVY KPVDLSKVTS KCGSLGNIHH KPGGGQVEVK SEKLDFKDRV QSKIGSLDNI 360
THVPGGGNKK IETHKLTFRE NAKAKTDHGA EIVYKSPVVS GDTSPRHLSN VSSTGSIDMV 420
DSPQLATLAD EVSASLAKQG L                                          441
```

**c**

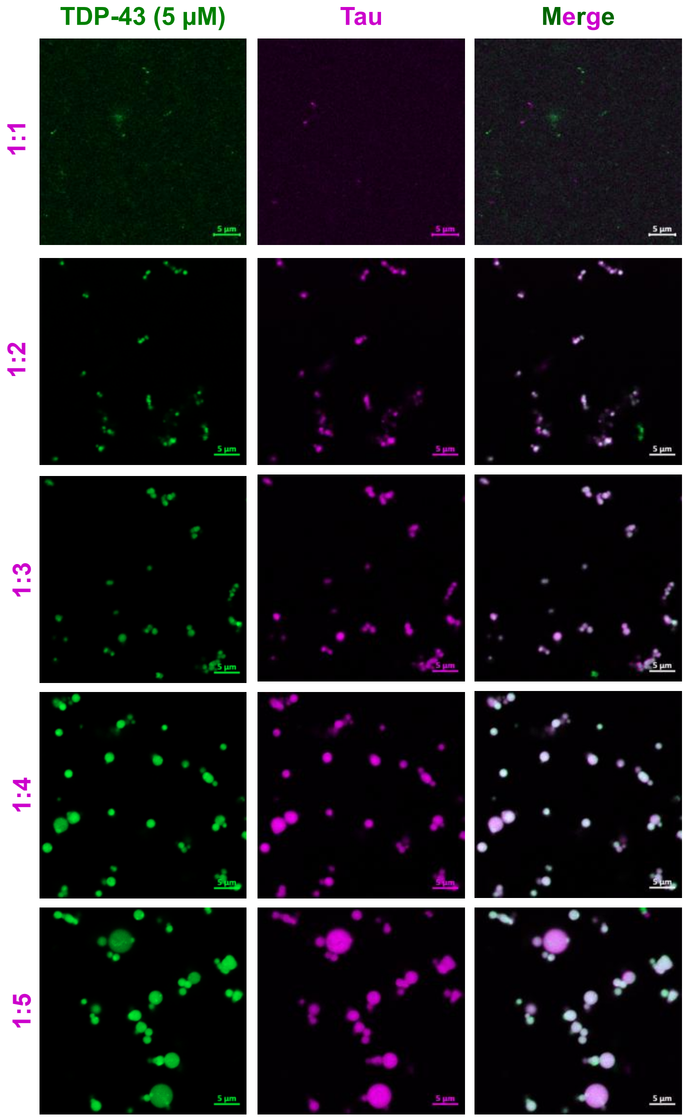


**Supplementary Figure 1:** Amino acid of TDP-43 (**a**) and tau (**b**), color-coded corresponding to the domain architecture shown in Fig. 1a,b. **c.** Representative two-color confocal Airyscan images with increasing ratios of tau (purple, 5-25 µM) with TDP-43 (green) fixed at 5 µM under our reaction conditions. Scale bar = 5 µm.

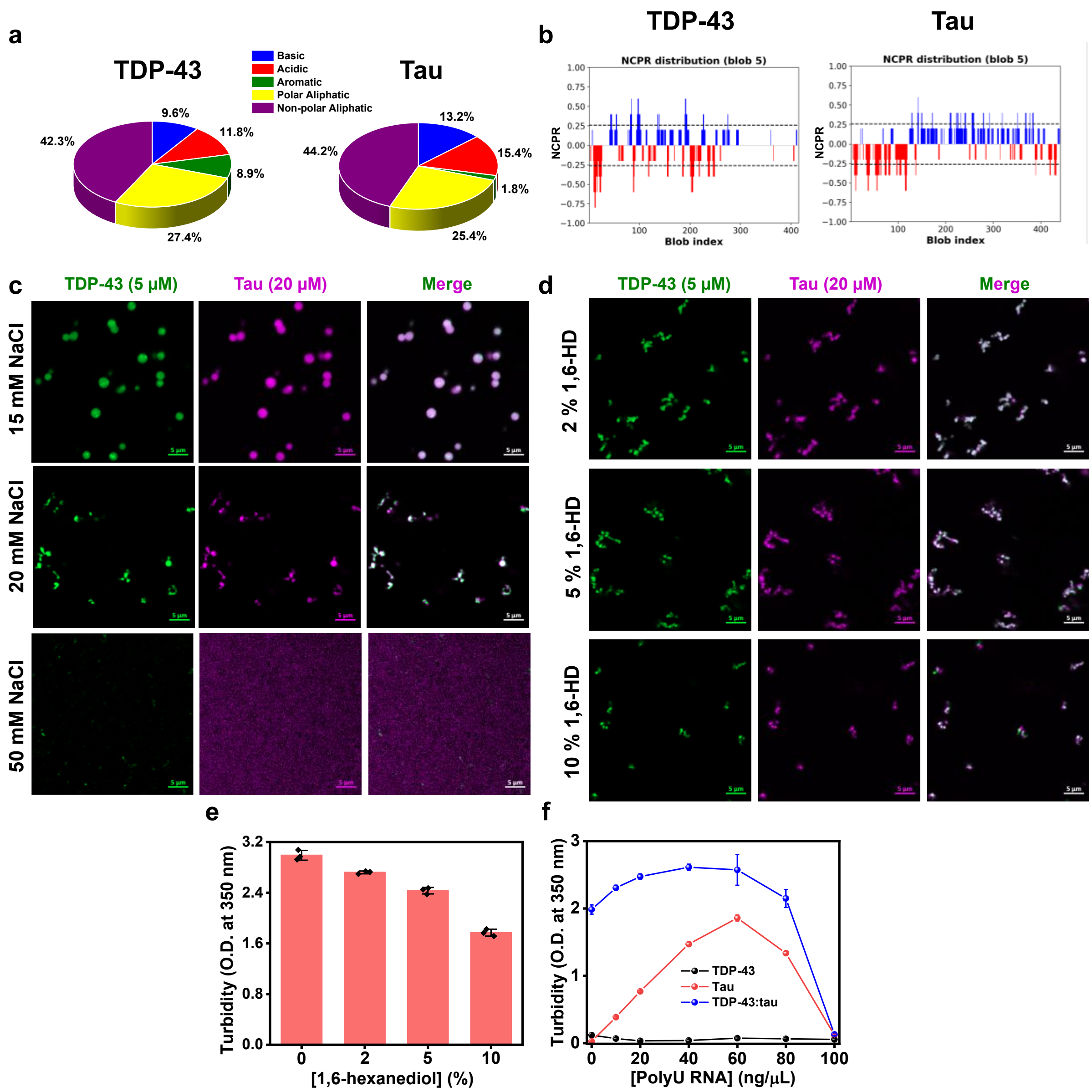


**Supplementary Figure 2: a.** A pie-chart representation showing the sequence composition of TDP-43 and tau, marking the proportion of various amino acid residues. **b.** Net-charge per residue (NCPR) plot created using CIDER analysis and calculated using a sliding window of 5 residues for TDP-43 and tau. Representative two-color confocal Airyscan images of TDP-43 (5 μM) and tau (20 μM) droplets formed in increasing concentration of NaCl (**c**) and hexanediol (**d**), depicting the effect of electrostatic and hydrophobic modulators on TDP-43:tau heterotypic condensation. Scale bar = 5 μm. **e.** Turbidity plot obtained for phase-separated reactions (5 μM TDP-43 and 20 μM tau) in the presence of increasing concentration of hexanediol. Data represent mean ± SD (n = 3 independent samples). **f.** Turbidity measurements for TDP-43 (black) and tau (red) individually and upon mixing at a 1:4 ratio (blue) in the presence of PolyU RNA, capturing the reentrant phase behavior of the heterotypic condensates over the concentration range of 0 ng/µL to 100 ng/µL RNA. Data represent mean ± SD (n = 3 independent samples).

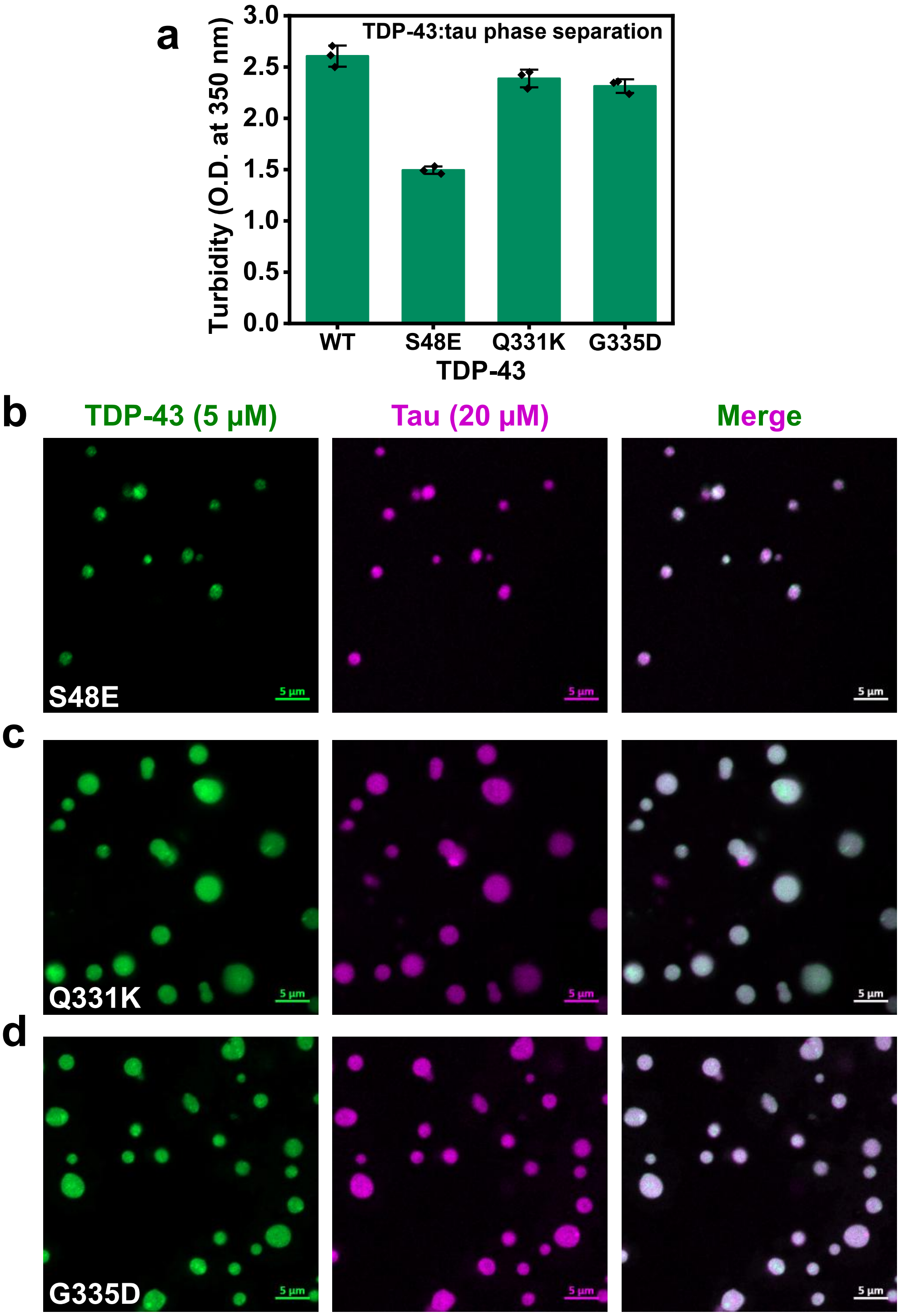


**Supplementary Figure 3: a.** Turbidity values measured at 350 nm for phase separation reactions upon mixing tau (20 µM) with wild-type or different TDP-43 mutants (5 µM). Data represent mean ± SD (n = 3 independent samples). Representative two-color confocal Airyscan images obtained for droplet reactions monitored in (**a**), showing formation of TDP-43:tau heterotypic droplets with TDP-43 S48E (**b**), TDP-43 Q331K (**c**), and TDP-43 G335D (**d**) in 20 mM HEPES, 15 mM NaCl, 1 mM DTT, pH 7.4. Scale bar = 5 µm.

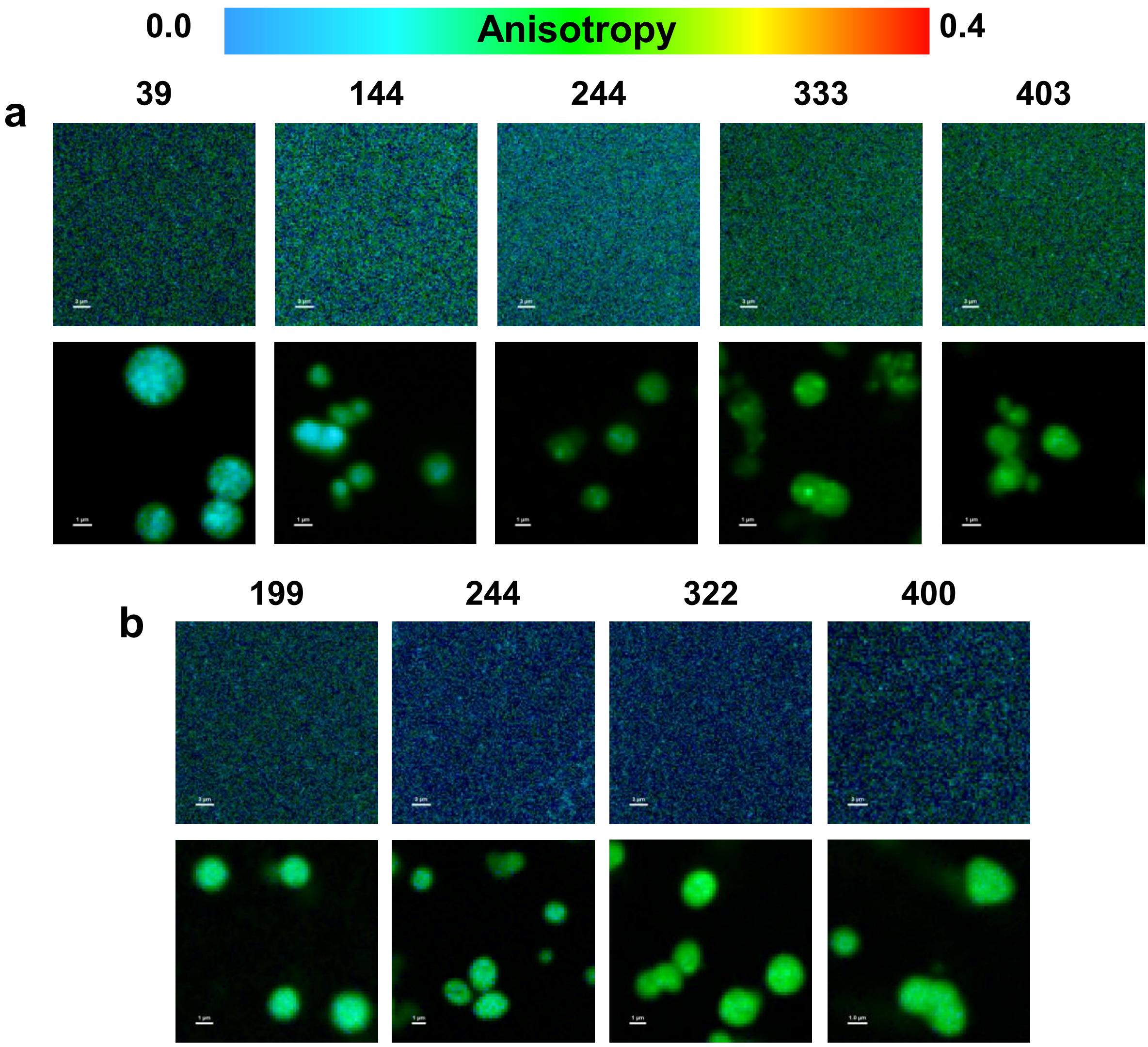


**Supplementary Figure 4:** Representative steady-state anisotropy images acquired for F5M-labeled single-cysteine variants of TDP-43 (**a**) and tau (**b**) in the dispersed and droplet phases. Scale bar = 3 µm and 1 µm for dispersed and droplet phase reactions, respectively. Anisotropy scale is shown at the top.

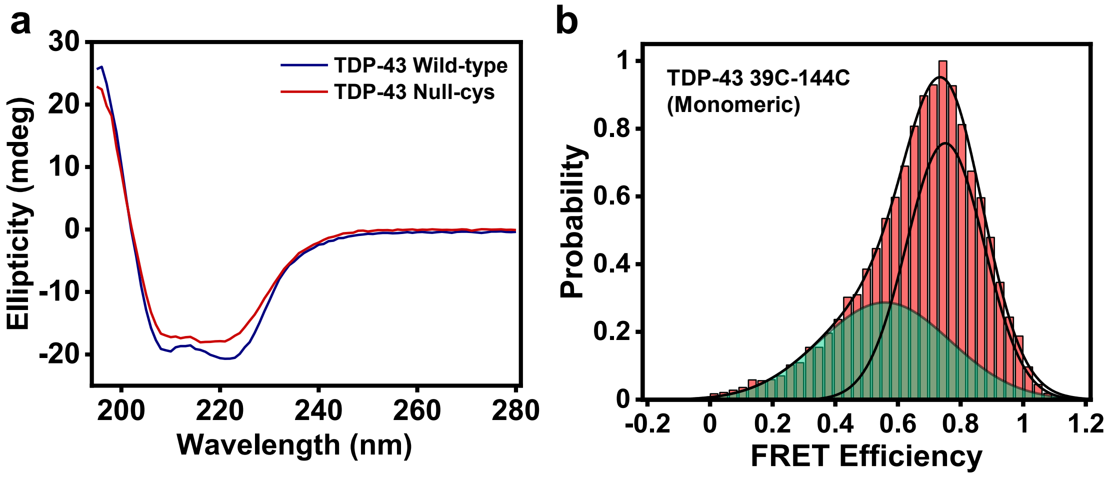


**Supplementary Figure 5: a.** Circular dichroism (CD) spectra comparing the secondary structural characteristics of MBP-tagged wild-type and null-cysteine TDP-43. **b.** Single-molecule FRET histograms obtained for monomeric TDP-43 39C-144C construct in the dispersed phase. The number of events at maxima was 1614, and the total number of events collected was > 20,000.

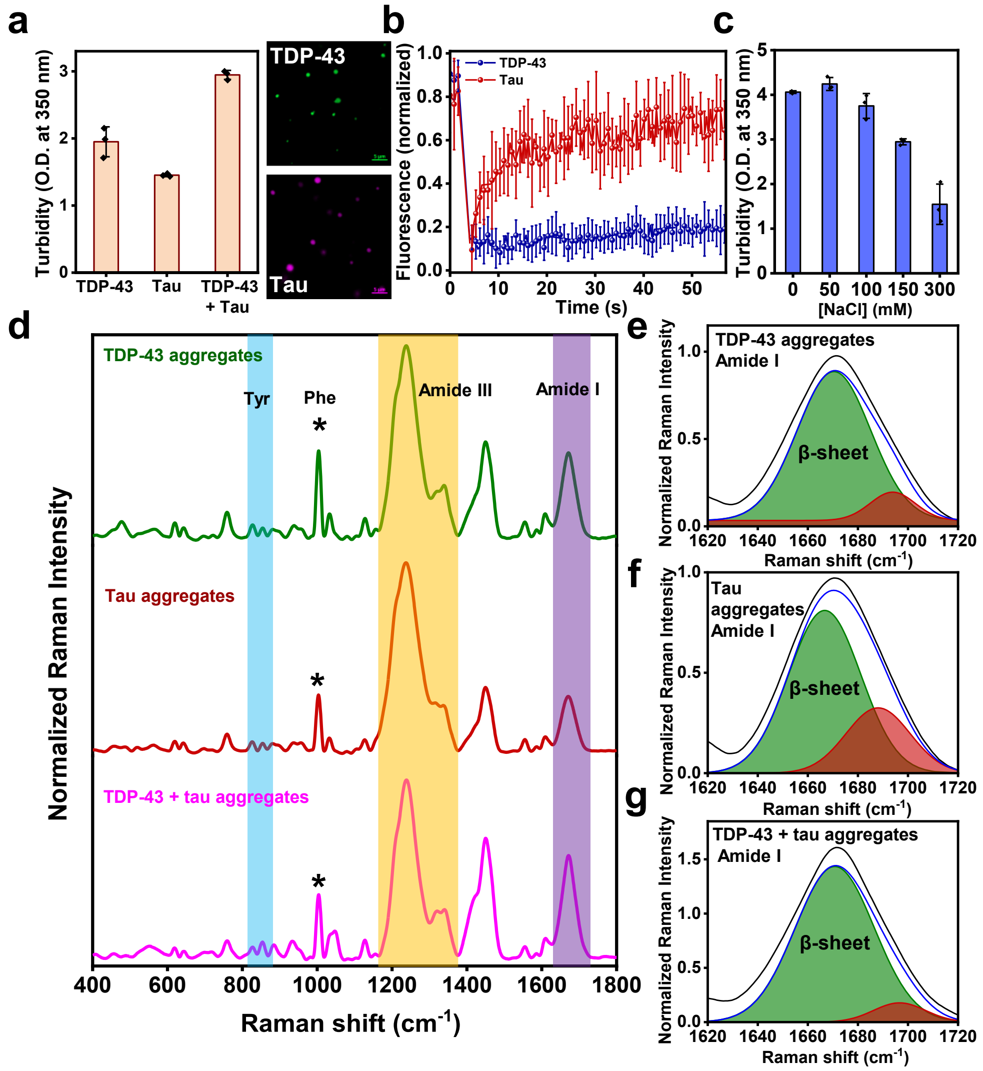


**Supplementary Figure 6: a.** Turbidity plot obtained for TDP-43 and tau homotypic and heterotypic phase separation in the presence of 10 % PEG and 150 mM NaCl. Data represent mean ± SD (n = 3 independent repeats). Corresponding images for TDP-43- and tau-only droplets are shown in the right panel. Scale bar = 5 µm. **b.** FRAP kinetics of both proteins within heterotypic condensates show a similar droplet interior to that observed for droplets formed under low-salt conditions. Data represent mean ± SD (n = 11 and 7 independent repeats for TDP-43 and tau, respectively). **c.** Salt-dependent turbidity plot of TDP-43:tau condensates confirming the role of electrostatic contacts underlying co-phase separation. Data represent mean ± SD (n = 3 independent repeats). **d.** Vibrational Raman scattering spectra obtained for aggregates formed by TDP-43 and tau alone and under co-phase separation conditions. The signature Raman bands, namely Tyrosine doublet, Amide I, and Amide III, are highlighted in the spectra. Data represent mean (n = 6 independent repeats). The Gaussian deconvolution obtained for the Amide I (**e-g**), highlighting the β-sheet (olive) and non-regular/disordered (red) contribution.

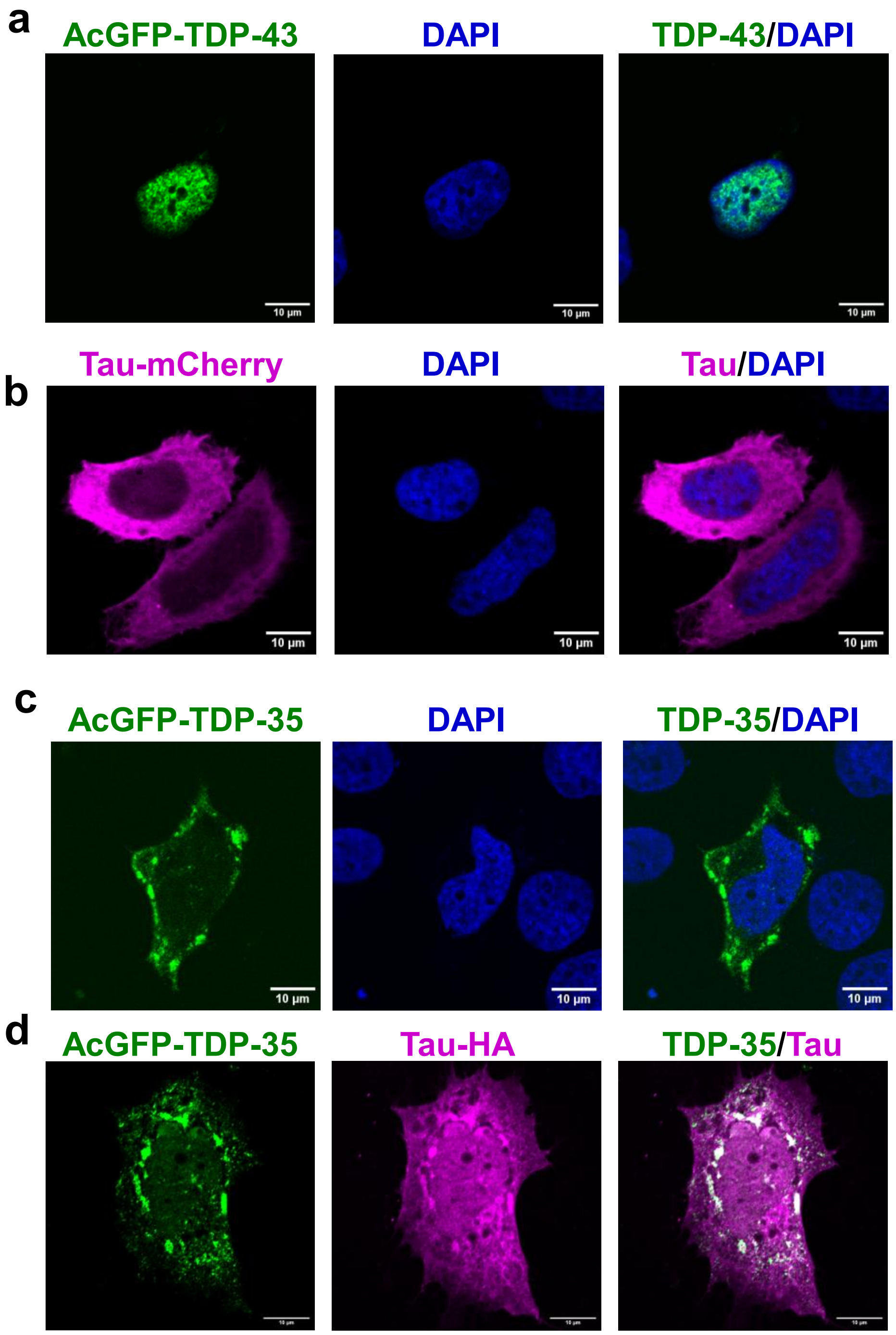


**Supplementary Figure 7:** Representative confocal images obtained for cells expressing AcGFP-TDP-43 (**a**), tau-mCherry (**b**), and AcGFP-TDP-35 (**c**) in HeLa cells subjected to 300 µM sodium arsenite treatment for 1 h. **d.** Representative confocal images showing partitioning of immunostained tau-HA with co-expressed AcGFP-TDP-35 under oxidative stress conditions. Scale bar = 10 µm for all images.

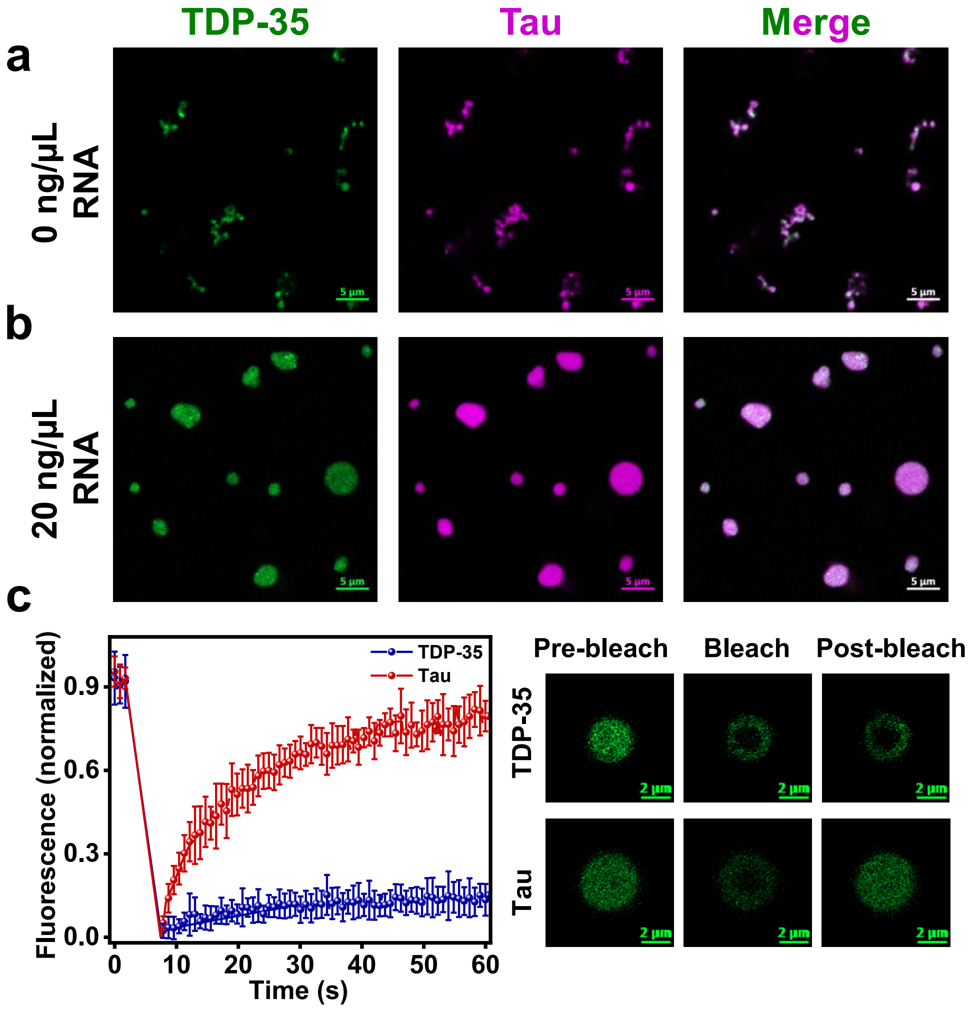


**Supplementary Figure 8:** Representative two-color Airyscan images showing heterotypic phase separation of recombinantly purified TDP-35 and tau in the absence (**a**) and presence (**b**) of 20 ng/µL PolyU RNA. Scale bar = 5 µm. **c.** FRAP kinetics obtained for A488-labeled TDP-35 (blue) and tau (red) within these RNA-induced droplets. Data represent mean ± SD (n = 10 and 8 independent repeats for TDP-35 and tau, respectively.

**Table S1.** List of constructs and the corresponding primers used in this study.

| Construct | Primer | Sequence (5' – 3') |
|---|---|---|
| TDP43 single cysteine(244C)-TEV-MBP-$His_6$-pJ4M | Forward | ATATATCATATGATGAGCGAGTATATTCGTG |
| | Reverse | ATATATCTCGAGCATGCCCCAACCGGA |
| TDP43 null cysteine-TEV-MBP-$His_6$-pJ4M | Forward | CAAATCGCGCAGAGCCTGAGTGGTGAGGATCTGATTATC |
| | Reverse | GATAATCAGATCCTCACCACTCAGGCTCTGCGCGATTTG |
| $His_6$-TEV-TDP-43 NRR (1-259) pJ411 | Forward | ATATATCATATGATGAGCGAGTATATTCGTG |
| | Reverse | ATATATCTCGAGCTAGTTAGAAATGTGAACGGAAATACC |
| $His_6$-TEV-TDP-35 (90-414) pJ411 | Forward | ATATATCATATGGCCAGCTCTGCGGTTAAAGTCAAGCG |
| | Reverse | ATATATCTCGAGCATGCCCCAACCGGA |
| TDP43 S39C-TEV-MBP-$His_6$-pJ4M | Forward | GCAGTTCCCAGGCGCCTGCGGCCTGCGCTATCG |
| | Reverse | CGATAGCGCAGGCCGCAGGCGCCTGGGAACTGC |
| TDP43 S144C-TEV-MBP-$His_6$-pJ4M | Forward | GATCTGAAAACCGGTCACTGCAAGGGTTTTGGCTTCG |
| | Reverse | CGAAGCCAAAACCCTTGCAGTGACCGGTTTTCAGATC |
| TDP43 S333C-TEV-MBP-$His_6$-pJ4M | Forward | CGCTGCAGTCTTGCTGGGGTATGA |
| | Reverse | TCATACCCCAGCAAGACTGCAGCG |
| TDP43 S403C-TEV-MBP-$His_6$-pJ4M | Forward | CGGCGGCTTTGGTTGCAGCATGGATA |
| | Reverse | TATCCATGCTGCAACCAAAGCCGCCG |
| | Forward | ATCGTAATCCGGTCGAGCAGTGCATGCGCGG |

| | | |
|---|---|---|
| TDP43 S48E-TEV-MBP-His$_6$-pJ4M | Reverse | CCGCGCATGCACTGCTCGACCGGATTACGAT |
| TDP43 Q331K-TEV-MBP-His$_6$-pJ4M | Forward | GCACAAGCGGCGCTGAAGTCTAGCTGGGGTATGATGG |
| | Reverse | CCATCATACCCCAGCTAGACTTCAGCGCCGCTTGTGC |
| TDP43 G335D-TEV-MBP-His$_6$-pJ4M | Forward | GCAGTCTAGCTGGGATATGATGGGCATGCTGGCGAGC |
| | Reverse | GCTCGCCAGCATGCCCATCATATCCCAGCTAGACTGC |
| TDP43 S333D-TEV-MBP-His$_6$-pJ4M | Forward | GGCGCTGCAGTCTGACTGGGGTATGATGG |
| | Reverse | CCATCATACCCCAGTCAGACTGCAGCGCC |
| TDP43 S342D-TEV-MBP-His$_6$-pJ4M | Forward | GCATGCTGGCGGACCAGCAGAACCAGT |
| | Reverse | ACTGGTTCTGCTGGTCCGCCAGCATGC |
| TDP43-pAcGFP-C1 | Forward | ATAGCCTCGAGCTATGTCTGAATATATTCGGGTAACCGAAGAT |
| | Reverse | ATAGCAAGCTTTTACATTCCCCAGCCAGAAGACTTAGAATC |
| TDP35-pAcGFP-C1 | Forward | ATAGCCTCGAGCTGCTTCATCAGCAGTGAAAGTGAAAAGAGC |
| | Reverse | ATAGCAAGCTTTTACATTCCCCAGCCAGAAGACTTAGAATC |
| Tau-pmCherry-N1 | Forward | ATATATATCTCGAGATGGCTGAGCCCCGCC |
| | Reverse | ATATATGGATCCGCCAAACCCTGCTTGGCCAGGGAGG |
| Tau-HA-pcDNA3.1(-) | Forward | ATATATATCTCGAGATGGCTGAGCCCCGCC |
| | Reverse | GGATCCCTATGCATAATCCGGAACATCATACGGATACAAACCCTGCTTGGCCAGGG |

| | | |
|---|---|---|
| Nh2htau (26-230)-pmCherry-N1 | Forward | ATATATCTCGAGATGCAGGGGGGCTACACCATGC |
| | Reverse | ATATATGGATCCGCACGGACCACTGCCACCTTCTTG |
| $Tau_{151\text{-}391}$-pmCherry-N1 | Forward | ATATATCTCGAGATGATCGCCACACCGCGGG |
| | Reverse | ATATATGGATCCGCTTCCGCCCCGTGGTCTGTC |

**Table S2.** Diffusion times obtained from FCS measurements for full-length and truncated variants of TDP-43 and tau in the dispersed and corresponding droplet phases.

| Construct | Dispersed/Droplet | Diffusion time (ms) |
|---|---|---|
| TDP-43 | Dispersed | 0.32 ± 0.07 |
| | TDP-43:tau droplet | 28.90 ± 12.40 |
| | TDP-43:tau droplet (20 ng/µL RNA) | 18.00 ± 5.51 |
| Tau | Dispersed | 0.98 ± 0.46 |
| | TDP-43:tau droplet | 9.09 ± 0.90 |
| | TDP-43:tau droplet (20 ng/µL RNA) | 6.72 ± 1.88 |
| TDP-43 NRR | Dispersed | 0.19 ± 0.05 |
| | TDP-43 NRR:tau droplet | 20.22 ± 3.70 |
| Tau | TDP-43 NRR:tau droplet | 18.85 ± 3.05 |
| TDP-43 | TDP-43:$tau_{151\text{-}391}$ droplet | 12.86 ± 3.63 |
| $Tau_{151\text{-}391}$ | Dispersed | 0.15 ± 0.04 |
| | TDP-43:$tau_{151\text{-}391}$ droplet | 7.18 ± 2.24 |
